# Lysis cassette-mediated exoprotein release in *Yersinia entomophaga* is controlled by a PhoB- like regulator

**DOI:** 10.1101/2023.02.08.527387

**Authors:** Marion Schoof, Maureen O’Callaghan, Charles Hefer, Travis R. Glare, Amber R. Paulson, Mark R.H. Hurst

## Abstract

Secretion of exoproteins is a key component of bacterial virulence and is tightly regulated in response to environmental stimuli and host-dependent signals. The entomopathogenic bacterium *Yersinia entomophaga* MH96 produces a wide range of exoproteins including its main virulence factor, the 2.46 MDa insecticidal Yen-Tc toxin complex. Previously, a high-throughput transposon-based screening assay identified the region of exoprotein release (YeRER) as essential to exoprotein release in MH96. The current study defines the role of the YeRER-associated <u>a</u>mbiguous holin/endolysin-based lysis <u>c</u>luster (ALC) and the novel RoeA regulator in the regulation and release of exoproteins in MH96. A mutation in the ALC region abolished exoprotein release and caused cell elongation, a phenotype able to be restored through *trans*-complementation with an intact ALC region. Endogenous ALC did not impact cell growth of the wild type, while artificial expression of an optimised ALC caused cell lysis. Using HolA-sfGFP and Rz1-sfGFP reporter, Rz1 expression was observed in all cells while HolA expression was limited to a small proportion of cells, which increased over time. Transcriptomic assessments found expression of the genes encoding the prominent exoproteins, including the Yen-Tc, was reduced in the *roeA* mutant and identified a 220 ncRNA of the YeRER intergenic that, when *trans* complemented in the wildtype, abolished exoprotein release. A model for *Y. entomophaga* mediated exoprotein regulation and release is proposed.

**Importance:** While theoretical models exist, there is not yet any empirical data that links ALC phage-like lysis cassettes with the release of large macro-molecular toxin complexes such as Yen-Tc in Gram-negative bacteria. In this study, we demonstrate that the novel *Y. entomophaga* RoeA activates the production of exoproteins (including Yen-Tc) and the ALC at the transcriptional level. The translation of the ALC holin is confined to a subpopulation of cells that then lyse over time, indicative of a complex hierarchical regulatory network. The presence of orthologous RoeA orthologue and a HolA like holin 5’ of an eCIS Afp element in *Pseudomonas chlororaphis* combined with the presented data suggests a shared mechanism is required for the release of some large macromolecular protein assemblies such as the Yen-Tc, and further supports classification of phage-like lysis clusters as type 10 secretion systems.

## INTRODUCTION

The release of exoproteins from bacteria plays a role in their nutrient acquisition, antimicrobial resistance, and delivery of toxins or other virulence factors enabling pathogenic bacteria to attach to and/or invade host tissues (1–3). To release exoproteins, several secretion pathways have evolved to transport proteolytic enzymes and other virulence factors across the protective lipid bilayer of the Gram-negative bacterial cell wall. These pathways range from simple protein channels and pores that span the cell wall to more complex multicomponent secretion systems, including the Type 1–9 secretion systems (T1-9SS) (4, 5).

In addition to cell wall-based secretion systems, indirect mechanisms such as the release of membrane vesicles and cell lysis via phage-like lysis cassettes (6–8) have been recognized as alternative mechanisms for protein release and/or transport (9–11). Palmer et al. (2021) modelled a T10SS on the *S. marcescens* ChiWXYZ-chitinase secretion pathway, where the unspecific holin pore ChiW allows translocation of proteins, including the hydrolase ChiX which degrades peptidoglycan in the cell wall (12). The ChiWXYZ can be described as a phage-derived lysis cassette, which upon activation causes the host cell to lyse (13).

The broad host range entomopathogen *Y. entomophaga* strain MH96 encodes multiple virulence factors including adhesins, Type 3 and Type 6 secretion systems, proteolytic enzymes, and insecticidal toxins (14). Many of these virulence genes are upregulated during haemocoelic infection of *Galleria mellonella* at 25°C (15). In Lauria Bertani (LB) broth at temperatures of ≤ 25°C *Y. entomophaga* releases large quantities of exoprotein including the Yen-Tc (16).The production of exoprotein is elevated through exponential growth and decreases through stationary phase (17). Based on reduced exoprotein in LB broth at 37°C, the production of these exoproteins including the Yen-Tc is likely under the control of a temperature sensor (16, 18).

Through use of a high-throughput secretion assay (HESA), a gene cluster termed *<u>Y</u>. <u>e</u>ntomophaga* <u>R</u>egion of <u>E</u>xoproteome <u>R</u>elease (YeRER) was identified which controls exoproteome production (including the Yen-Tc) by this species (17). The YeRER comprises a predicted phage-like lysis cassette designated the <u>a</u>mbiguous holin/endolysin-based lysis <u>c</u>luster (ALC) – encoded by *holA, pepB, rz,* and *rz1-* and a predicted DNA-binding regulator (RoeA) separated by a 1086 bp intergenic region. The HESA identified three transposon insertions (H45, H4 and H31) respectively located 70, 120 and 237 nucleotides 5’ of the ALC *hol*A initiation codon, respectively, and one transposon mutant (H12) with a single transposon insertion within *roeA*, which significantly reduced exoprotein production. Further to this, a 3-bp deletion of a spontaneous exoprotein deficient mutant strain K18 was identified in the YeRER intergenic region 131 bp 5’ of the *roeA* initiation codon (17). Based on these findings it is likely that RoeA is a global activator of MH96 exoprotein production via control of the lysis cassette.

*In silico* analysis of the 15 kDa RoeA protein identified the protein to be of the PhoB family of two-component regulators (TCR) (19, 20). In this context PhoB-like proteins comprise a membrane-bound histidine kinase domain that responds to specific environmental stimuli, and a receiver domain which acts as a transcriptional regulator upon its phosphorylation by its cognate histidine kinase (21–24).

To date the *Yersinia enterocolitica* W22703 thermoregulatory mechanism involving the LysR type regulators tcaR1 and tcaR2 located 5’ of the W22703-encoded *tc* genes and the Tc-PAI*_Ye_* lysis cassette juxtaposition to TcaC and TccC are the most studied example of regulation and Tc release (25, 26). In this system it has been proposed the lysis cassette, comprised of *holY, elyY, yRz* and *rRz1,* can cause cell lysis, but no biological proof of the function has yet been demonstrated.

Based on the unique properties of RoeA, its associated ALC lysis cassette and its role in exoprotein production, we have used a combination of fluorescent and enzymatic reporter genes, microscopy, and transcriptomics to determine the mechanism by which MH96 lysis cassette-mediated exoprotein release is regulated.

## RESULTS

### Exoprotein release by the ambiguous lysis cluster (ALC) of the Yersinia entomophaga region of exoprotein release (YeRER)

Based on the importance of the YeRER in exoprotein production we undertook gene synteny analysis of YeRER components to genomes in the current database. Genome comparison identified the location of other lysis-cassettes within a toxin complex (Tc) of the *Yersinia* genus (Fig. 1A). The composition of these lysis cassettes is typified by the presence of a holin, a peptidase and one to two spanin proteins, in which the MH96 ALC resembles a classical phage λ lysis cassette comprising a holin, an endolysin and spanin complex Rz/Rz1 (Fig. 1A, Fig. S1A). While cell lysis of phage λ is typically controlled by a dual translational start in its λ holin gene S, forming the holin (S105)- antiholin (S107) complex (27, 28), the holin (HolA) of the *Y. endomophaga* MH96 lysis cassette does not contain a dual-start motif, and therefore does not form an anti-holin to regulate cell lysis. Phylogenetic assessment of HolA with selected orthologues of the *Enterobacteriaceae* shows that closely related holins are also devoid of a HolA dual start motif (Fig. S1). Furthermore, gene arrangements differed between the strains, but unique to *Y. entomophaga* and *Y. nurmii* are the overlapping *holA* and *pepB* open reading frames (Fig. S1B).

**Figure 1:**
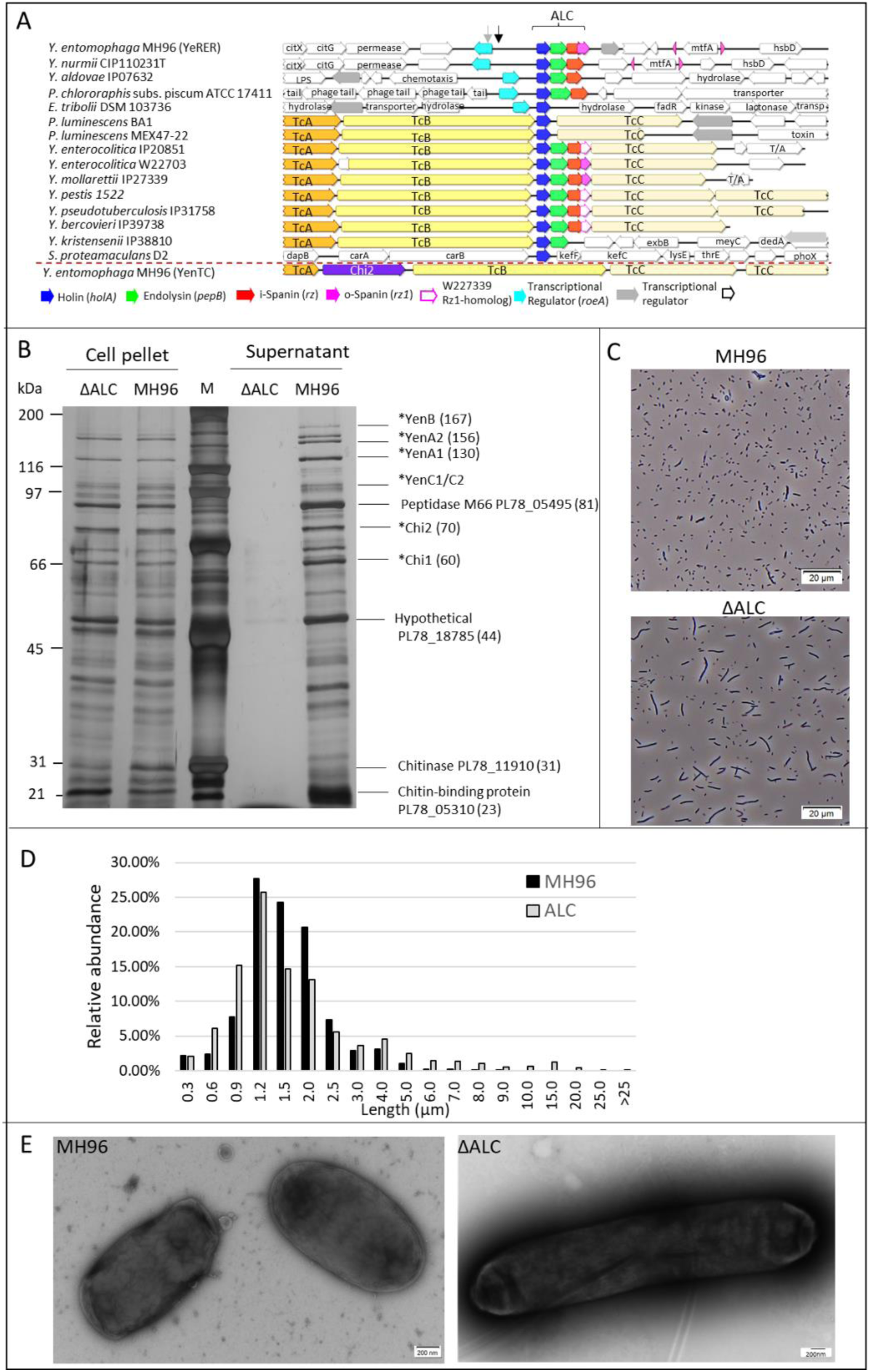
A) Gene synteny of the ALC in the YeRER to selected *Yersinia* sp., *Serratia proteamaculans*, *Photorhabdus luminescens, Pseudomonas chlororaphis* and *Enterobacillus tribolii,* strain identifiers are indicated. Associated Tc clusters are cross referenced to the *Yersinia entomophaga* MH96 Yen-Tc cluster positioned below red dashed line. Gray and black arrows denote the position of the respective H12 and K18 mutations. B) SDS-PAGE (10%) of culture supernatant from strains MH96 and ΔALC. C) Light microscopy of MH96 and ΔALC, in which some ΔALC cells are elongated. Scale bar of 20 µm indicated. D) Distribution in relative abundance of cell lengths in cultures of MH96 and ΔALC derived from light microscopy images. Total number of cells measured – 6681 per culture. E) Negative stain of MH96 and ΔALC using transmission electron microscopy, confirming cell elongation in ΔALC.

Based on phylogenetic assessment, the MH96 HolA is most closely related to the *Y. nurmii* HolA and together forms a cluster with *Serratia proteamaculans, Enterobacillus tribolii, Yersinia aldovae*, and *Photorhabdus luminescens* which are distinct from holins of the other *Yersinia* species, such as *Y. enterocolitica* (Fig. S2A). Similarly, the *Y. entomophaga* MH96 M15 endopeptidase, which is absent in the *P. luminescens* associated Tc clusters, is closely related to *Y. nurmii*, but less closely related to other *Yersinia* species (Fig. S2B).

To validate the role of the ALC in cell lysis and exoprotein release, we constructed an ALC deletion mutant, ΔALC (Table S1). Initially we quantified cell viability of the ALC mutant using viable colony forming unit (CFU) counts and live cell counts using microscopy of LIVE/DEAD stained cells of MH96 and ΔALC through the growth phase. Neither live cell counts nor CFU counts (Fig. S3) revealed a significant change in cell viability in cultures of MH96 and ΔALC which either suggests that the ALC does not cause cell lysis, or alternatively that ALC-mediated cell lysis and death is compensated for by growth of the non-lysed cell population and is therefore not detectable through CFU counts.

While the deletion of the ALC did not significantly alter cell counts, the deletion reduced the release of exoprotein including the Yen-Tc and several prominent bands identified by SDS-PAGE which are present in MH96. LC-ESI-MS/MS of five of these prominent MH96 protein bands identified a haemolysin (PL78_09600), a glycosyl hydrolase family protein (PL78_11910), a chitin-binding protein (PL78_05310), a peptidase M66 (PL78_05495) and a protein of unknown function (PL78_18785) (Fig. 1B, Fig. S4, Fig. S5).

Light-microscopy of ΔALC cells at early stationary phase (9.4 log_10_ CFU mL^-1^) (Fig. 1C), when exoprotein release was highest in MH96, revealed a proportion (>13%) of ΔALC cells are elongated with cells up to 37 µm in length observed, compared to average size of 1.5 ±0.6 µm of MH96 (Fig. 1D). Under electron microscopy, the ΔALC cell walls were continuous and non-septate (Fig. 1E).

### Induction of an artificial ALC with non-overlapping reading frames causes rapid cell lysis

To demonstrate the potential role of the ALC in cell lysis of MH96, the arabinose inducible vector pAY-ALCΔ*rz1* harbouring the ALC with its native gene arrangement of *holA, pepB* and *rz* but devoid of *rz1* (Fig. 2A) was induced and no change in optical cell density (OD_600_) in either MH96 (Fig. 2B) or *Escherichia coli* DH10B (Fig. 2C) was detected. This likely reflects the requirement for back translational coupling (termination-reinitiation), which is typical for λ phage-like lysis (29, 30). To uncouple the ALC, the MH96 ALC-encoding ORFs were separated (ALC-opt) resulting in the vectors pAY-ALC-opt, pAY-ALCΔ*pepB*-opt, pAY-ALCΔ*rz*-opt and pAY-ALCΔ*rz1*-opt (Fig. 2A; Table S2). Subsequent induction of pAY-ALC-opt resulted in rapid cell lysis and a concurrent decrease of OD_600_ from 1 to 0.2 within 15 min post induction (mpi) (Fig. 2D, Movie S1). Live-cell imaging of pAY-ALCΔRz1-opt revealed cell elongation and localised bulging of cells leading to membrane blebbing (15 mpi) prior to cell lysis (20 mpi) (Fig. 2D, Movie S2). In some cells, typically in elongated cells, one to three small localized dark areas were observed (Fig. 2D), which disappeared within 15 min. During the cell elongation, brightening at cell poles was observed (10 mpi). In MH96, the expression of the ALC missing either i-spanin Rz or o-spanin Rz1 in pAY-ALCΔ*rz*-opt and pAY-ALCΔ*rz1*-opt, caused rapid cell lysis as seen in ALC-opt expression (Fig. 2B). However, lysis activity of the ALC-opt in *E. coli* increased in the absence of either spanin (Fig. 2C), which may reflect cell wall composition differences between these species. In absence of *pepB,* slow cell lysis was observed when inducing pAY-ALCΔpepB-opt in *E. coli* (Fig. 2C).

**Figure 2:**
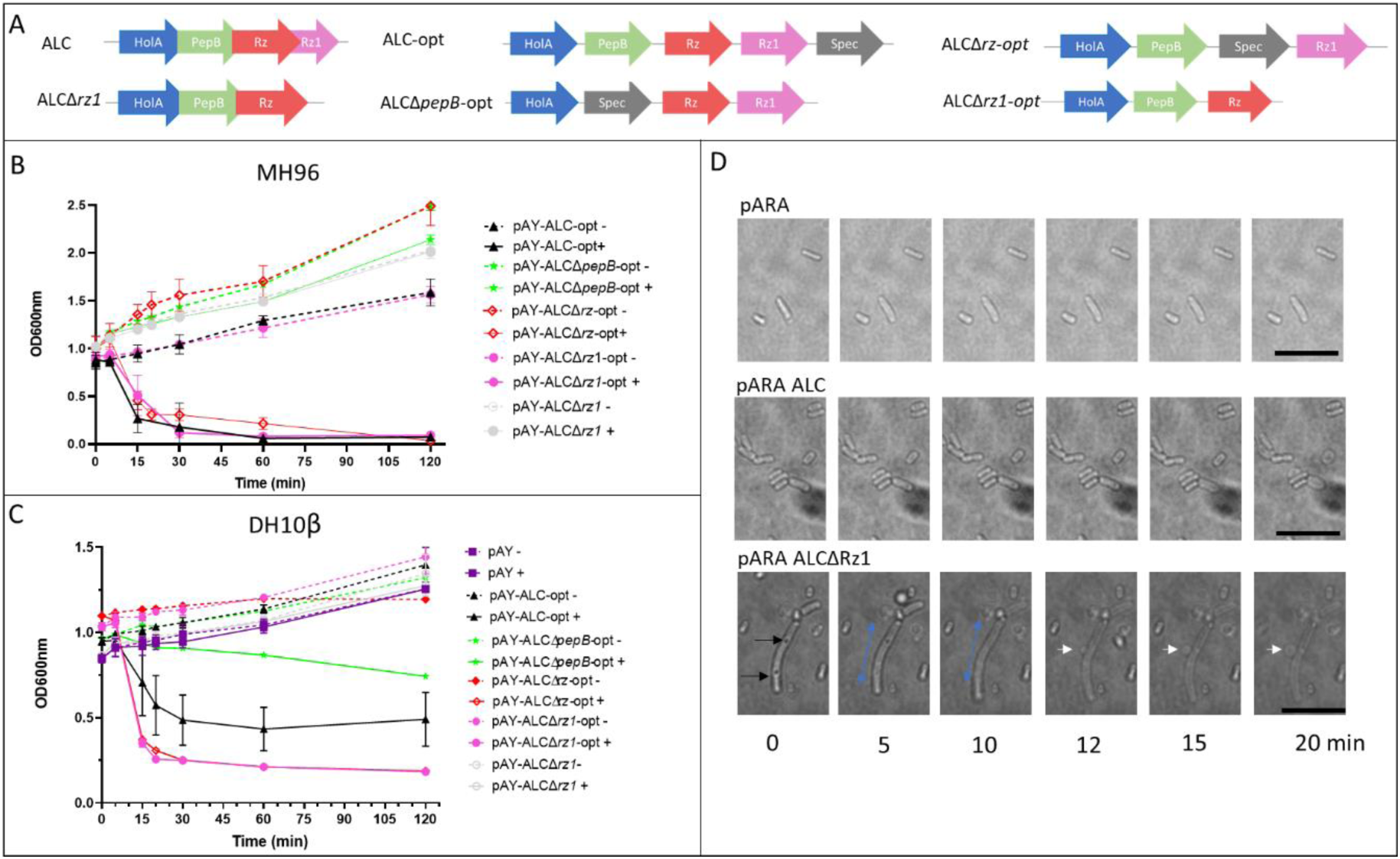
OD_600_ assessment of lysis activity of the pAY-ALC constructs induced with 0.02% arabinose (+) compared to uninduced culture (-) over 120 minutes. A) MH96 or B) *E. coli*; C) Live-cell imaging of cells upon induction with 0.6% arabinose, over 20 minutes. Black arrows denote localized dark areas, blue arrows denote directional cell elongation, white arrows denote membrane blebbing, scale bar: 10 µm. D) Schematic of pAY-ALC lysis variant (refer Table S2 for synthesised DNA and Table S3 vector details).

### Cell lysis is restricted to a subset of cells expressing HolA

To correlate ALC activity with MH96 cell lysis, the translational HolA-sfGFP and Rz1-sfGFP constructs were assessed in LB broth cultures at 25°C. During early exponential growth (6-8 h), 15 % of HolA-sfGFP cells were fluorescing, increasing to 44% at mid-late exponential phase (10-12 h post inoculation (hpi)) and decreasing slightly to 41% at stationary phase (>12 hpi). The HolA*-*sfGFP signal was observed over the entire cell membrane (Fig. 3) and in elongated cells through these growth phases. Parallel assessments of Rz1-sfGFP found that all stationary phase cells were fluorescing with the entire cell membrane fluorescent (Fig. 3).

**Figure 3:**
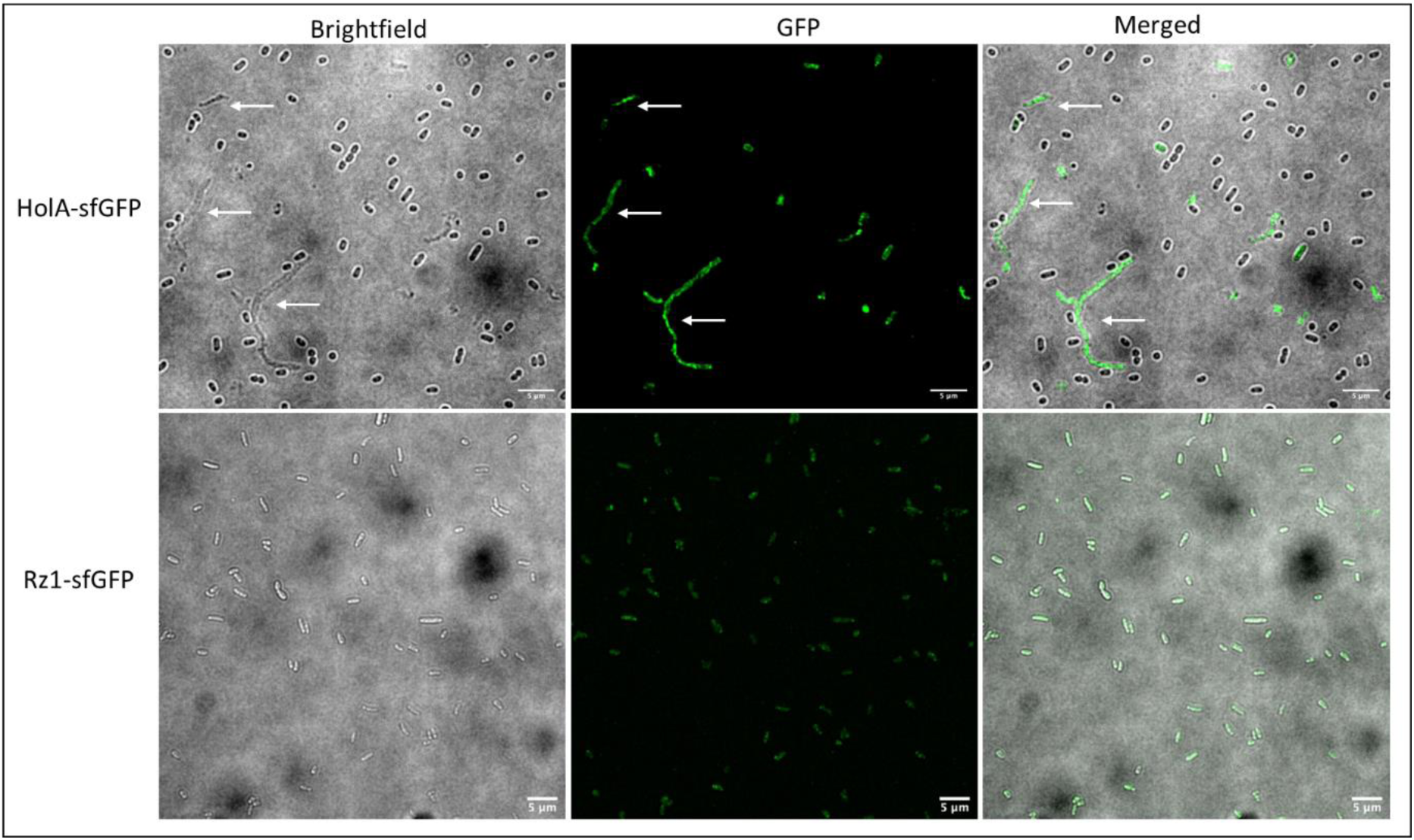
Light and fluorescent microscopy of MH96 cells expressing HolA-sfGFP and Rz1- sfGFP. Merged image shown. White arrows denote elongated cells. Scale bar (5µm) indicated.

### The PhoB-like regulator RoeA alters ALC, exoprotein and global gene expression

As previously noted, a HESA transposon insertion H12 in *roeA* abolished exoprotein which based on proximity to the ALC and gene synteny suggests that RoeA may regulate the ALC.

Through amino acid alignments and correlation of RoeA to the resolved structures of the TCR’s, CadC (31) and OmpR (20, 32), RoeA shares a PhoB-like helix-turn-helix (HTH) motif, comprising 3 α-helices, of which α2 and α3 are connected by a DNA loop forming the DNA binding structure (33, 34) in which the DNA binding motif resides (Fig. 4). Of interest, the C-termini of RoeA and RoeA-like proteins extend up to 40 amino acids compared to PhoB-like proteins, with the exception of CadC, which harbours a N-terminal HTH motif. Unlike PhoB, RoeA has no cognate phosphorylation domain.

**Figure 4:**
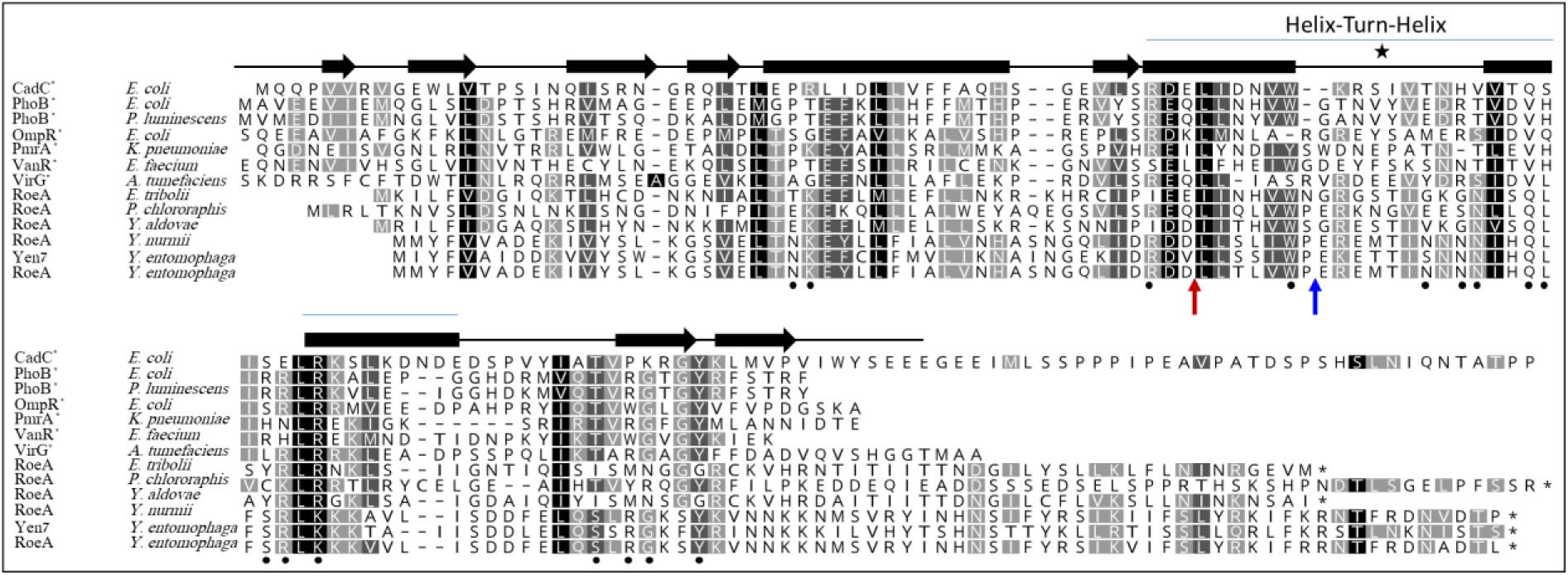
Amino acid alignment and secondary structure prediction of RoeA and its Yen7 orthologue and selected PhoB type regulators. Black filled circles denote PhoB residues linked to DNA binding (33) and area marked by a star denotes HTH-motif (34). Red and blue arrows denote respective MH96ΔroeA151::Spec and H12 mutation sites.

Assessing the non-redundant National Center for Biotechnology Information’s (nrNCBI) protein database, RoeA orthologues were identified in a limited number of bacteria, mainly within Yersiniaceae (sharing 30-100% amino acid identity) and Enterobacteriaceae (<30% amino acid identity) (Fig. S6). Of note, a RoeA orthologue (40% amino acid similarity to RoeA) is located 5’ of the Afp/ PVC-like eCIS protein complexes of *Pseudomonas chlororaphis* and is in juxtaposition to a ALC HolA orthologue (Fig. 1A). Interestingly, a second MH96 RoeA-homolog Yen7 is located 5’ of Yen-Tc, which is a similar position to the LysR-like regulator tcaR of other Tc-encoding *Yersinia* strains (12) .

Attempts to delete *roeA* in its entirety were unsuccessful, while SDS-PAGE assessment of MH96Δ*roeA151::Spec* containing a spectinomycin cassette 151 bp 3’ of the *roeA* initiation codon (Table S1) showed an exoproteome profile similar to MH96 (data not shown). This contrasts to the highly reduced exoprotein profile of the transposon insertion H12. Through the positioning of these mutations on the RoeA amino acid sequence (Fig. 4), the H12 insertion prevents translation of the entire RoeA HTH-motif, while the MH96Δ*roeA151::Spec* insertion retained 90% of the α2 helix and a partial HTH motif which may enable its functionality. Based on the different exoproteome profile of H12 and MH96Δ*roeA151::Spec* the genome of H12 was sequenced from where a single transposon insertion within *roeA* was validated and the H12 *roeA*^-^ strain was used in subsequent assessments.

To further define the role of RoeA in the transcriptional regulation of exoproteins and their release mechanism, the *roeA* H12 mutant and the wild-type MH96 were subjected to transcriptomic assessments, targeting early stationary growth phase at approximating 9.6. log_10_ CFU mL^-1^. Through DEseq2 analysis of the transcriptome, the H12 *roeA* mutant resulted in the differential expression of 2235 (53%) genes relative to MH96. For analysis purposes we concentrated on genes with transcription levels increased or decreased by log_2_ fold >|1| and identified 406 genes that were significantly overexpressed and 500 significantly down-regulated (Fig. 5).

**Figure 5:**
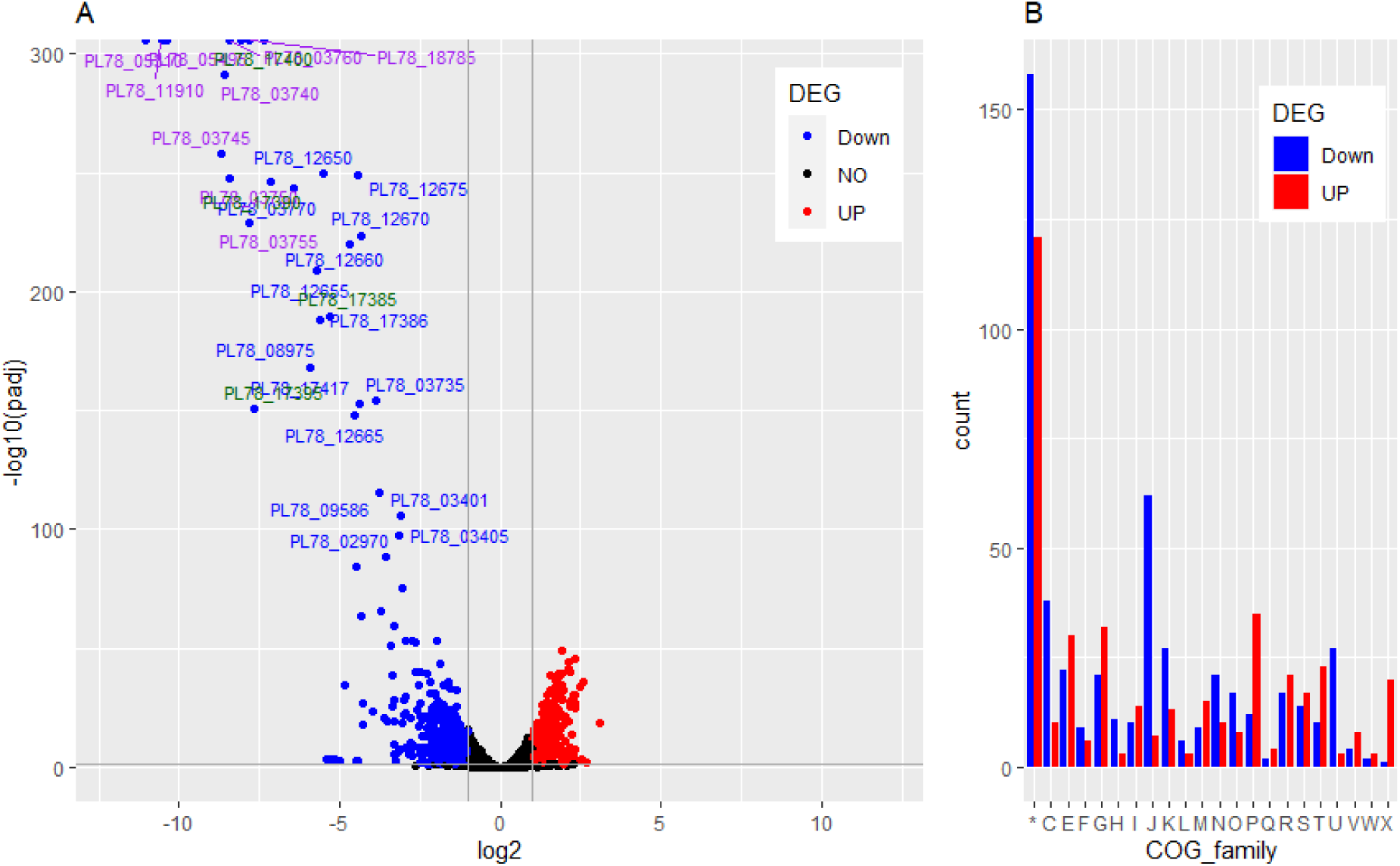
DeSeq2 analysis of gene expression levels of genes that are significantly regulated with p_adjust_< 0.005 in the *roeA* mutant H12 compared to MH96. A) Volcano plot of down-regulated genes in H12 by a fold change of log_2_ <-2 (blue) and upregulated genes by a fold change of log_2_ >2 (red). Genes expressing exoproteins are marked in purple, and YeRER associated genes are marked in green. Genes with fold change log2 between −2 and 2 are indicated in black. B) Distribution of COG classification of the differentially described genes sorted by up- and down-regulated genes. * denotes genes that had not been assigned to a COG-classification: C, Energy production and conversion; E Amino acid metabolites and transport; F Nucleotide metabolism and transport; G, Carbohydrate metabolism and transport; H, Coenzyme metabolites; I, Lipid metabolism; J, Translation; K, Transcription; L, Replication and repair; M, Cell wall/membrane/envelop biogenesis; N, Cell motility; O, Post-translational modification, protein turnover, chaperone functions; P, Inorganic ion transport and metabolism; Q, Secondary Structure; T, Signal Transduction; U, Intracellular trafficking and secretion; V, Defense mechanisms, W, Extracellular structures; X, Mobilome: Prophages, Transposons, R, General Functional Prediction only; S, Function unknown.

While most differentially regulated genes in *roeA* mutants have no assigned COG-classification, the H12 *roeA* mutation affected genes with a wide range of functions, with greater effects on genes involved in: i) post-translational modification, such as heat-shock proteins, DnaJK and GroEL molecular chaperones; ii) translation, such as ribosomal proteins; and iii) intracellular trafficking and secretion, such as genes of the T2SS (Fig. 5B). Other genes function in energy production, and carbohydrate, amino acid and inorganic ion transport and metabolism (Fig. 5B, File S1). Importantly the transcription of genes encoding for the ALC, its associated *roeA,* Yen-Tc, the *roeA* homologue *yen7* and the genes encoding the predominant wildtype MH96 exoproteins PL78_18785, PL78_05495, PL78_11910, and PL78_05310 (validated through LC-ESI-MS/MS) were significantly reduced in the *roeA* H12 mutant (File S1).

### Temperature dependent regulation of RoeA and its effect on exoprotein production

Based on the role of RoeA in the transcription of exoproteins including the Yen-Tc and the effect of temperature on virulence regulation *in vitro* (35) and *in vivo* in *G. mellonella* at 25°C and 37°C (15), we investigated the effect of temperature on the translation of RoeA and the Yen-Tc associated Chi1, as an exoprotein proxy.

Using a P*_roeA_::lacZ cis-*merodiploid strain cultured at 25°C in LB broth, the β-galactosidase levels increased during exponential growth phase but stabilised at stationary growth phase (Fig. 6), which correlates with the increased exoprotein levels produced through the exponential growth phase (17). In parallel assessments at 37°C, P*_roeA_::lacZ* β-galactosidase levels were significantly reduced (Fig. 6). At 37°C the β-galactosidase activity of P*_chi1_::lacZ* grown at 37°C was significantly reduced compared to 25°C (Fig. 6).

**Figure 6:**
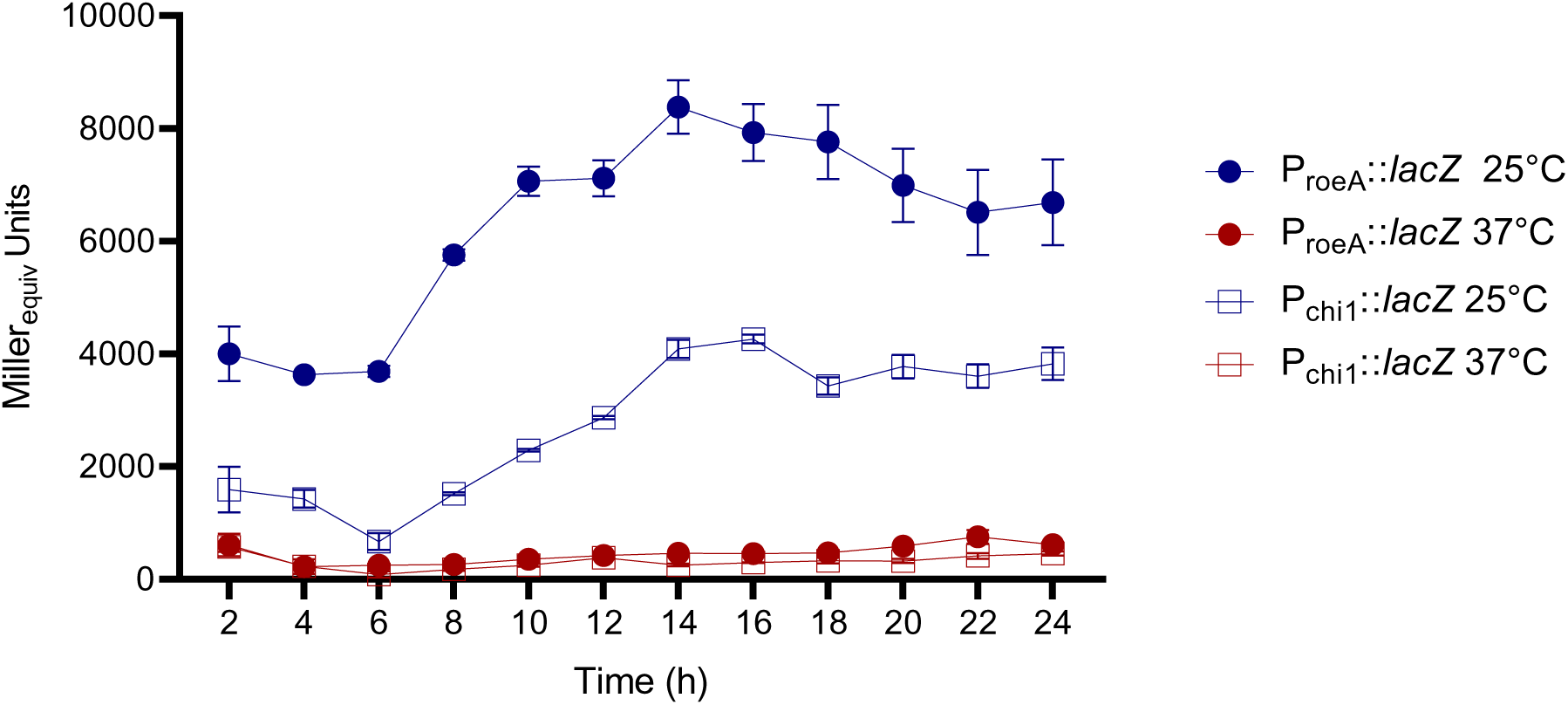
β-galactosidase production of MH96 *cis-*merodiploid P*_chi1_*::*lacZ* and P*_roeA_*::*lacZ* strains at 25°C and 37°C over a 4-24 h duration. Error bars represent ± SD of biological duplicates and technical replicates from the same culture flask.

### The YeRER intergenic region affects the regulation of the ALC and RoeA

As previously demonstrated in the HESA, three mutations (H4, H31, H45, 5’ of the ACL) and the spontaneous K18 mutation (5’ *roeA*) were identified in the YeRER intergenic region (17), suggesting an important role for the intergenic region in ALC regulation and therefore exoprotein release. Through transcriptome assessments, the predicted transcriptional start site of *roeA* and *holA* was located 3 bp and 350 bp 5’ of the respective gene (Fig. S7A). Bioinformatic analysis of the 350 bp region identified a 153 bp ORF (termed ORF153) including a predicted ribosomal binding site (5’ GGA 3’) 11 bp 5’ of the initiation codon of ORF153. The translated product of ORF153 had no similarity to proteins in the current nrNCBI protein sequence database (30/10/2022).

A saliant finding through the assessments of the RNAseq mRNA reads was the identification of a low number of mRNA reads spanning a 220bp region of the YeRER intergenic region in both WT MH96 and the *roeA-*H12 mutant, and the predicted ncRNA designated ncALC220 (Fig. S7A). Based on this information we further interrogated the YeRER nucleotide sequence for potential regulatory signatures (Fig. S7B). The YeRER intergenic region is AT rich with 33.7 % G+C relative to 48.6% G+C of the MH96 genome and harbours several protein binding motifs including those for PhoB and Fur-like proteins identified using the Prodoric software (Table S4). The *Vibrio cholerae* cyclic AMP receptor protein (CRP) DNA binding motif was found in the core sequence of the degenerate repeats (Fig. S7B). Further investigation of the nucleotide sequences 5’ of *roeA* and the *roeA* homologue *yen7*, identified the same H-NS binding site (36) 32 bp 5’ of their respective initiation codons (Fig. S7C). Additionally, Hfq binding motifs (5 AATAATA 3) (34) can be found in the intergenic region including one in the ncALC220 (Fig. S7B).

Based on the bioinformatic analysis of the YeRER associated intergenic region and the potential role of ncALC220 in YeRER regulation, a series of pACYC184 (p184) based vectors (Fig. 7A, Table S3) were constructed enabling their effect on exoprotein production in either a MH96, ΔALC, H4, or K18 background to be determined. The vectors comprise various sections of the intergenic region, ncALC220, and the intergenic region where the ncALC220 sequence was deleted (p184INTΔncALC220) (Fig. 7).

**Figure 7:**
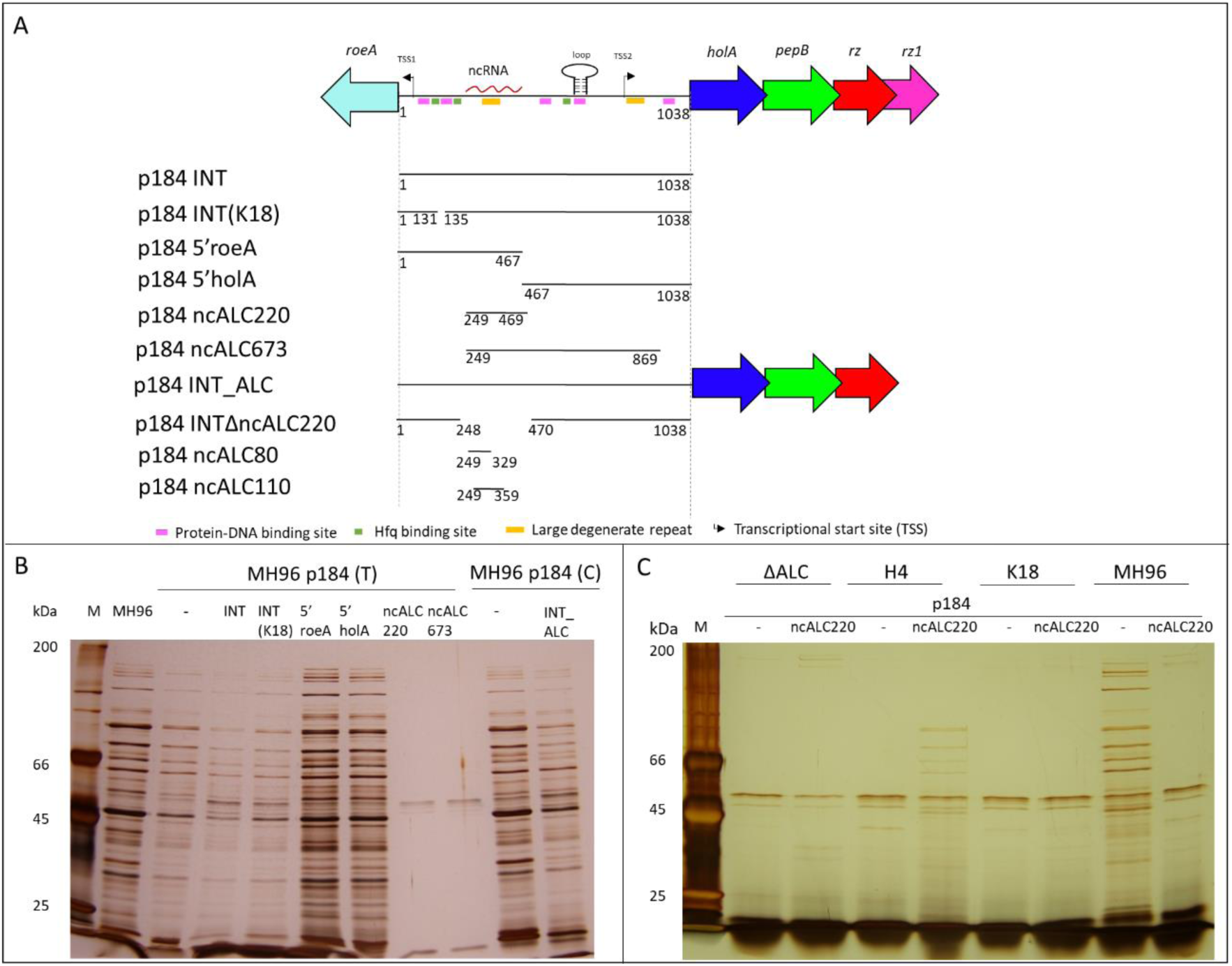
A) Schematic of the YeRER, including putative secondary structures and transcription factor binding sites. Depicted are the *trans* complementation vector series p184 p184INT, p184 INT(K18), p184 5’roeA, p1845’holA, p184ncALC220, p184ncALC673, p184INT_ALC, p184INTΔncALC220, p184ncALC80, and p184ncALC110 used for *trans* complementation. B) SDS-PAGE (10%) of the exoproteome profiles from culture supernatant of MH96 *trans* complemented with the p184 vector series each grown in the presence of 10 µg/mL tetracycline (T) (p184 T), empty vector (p184 -), and p184INT_ALC grown in the presence of 30 µg/mL chloramphenicol (C) (p184 C) (empty vector (p184 -). C) SDS-PAGE (10%) of the exoproteome profiles from supernatant of MH96, K18, H4 and ΔALC containing empty vector pACYC184 (p184 -) or pACYC184 containing the ncALC220 (p184 ncALC220), each grown in the presence of 10 µg/mL tetracycline.

Through assessments by SDS-PAGE, the *trans* complementation with either p184INT or p184ncALC220 in MH96, resulted in reduced exoprotein relative to the vector only MH96 control (Fig. 7B). Importantly the same intergenic region devoid of the 220 bp ncRNA (p184INTΔncALC220) did not alter the MH96 exoprotein profile (Fig. S8A). Truncated versions of ncALC220, namely ncALC80 (1-80 bp of ncRNA220) and ncALC110 (1-110 bp of ncRNA220), did not alter exoprotein production or cell morphology when *trans*-complemented in MH96 (Fig. S8A). The *trans* complementation of p184ncALC220 with either ΔALC or K18 did not restore exoprotein production (Fig. 7C), however a partial restoration was observed in H4 (transposon insertion 5’ of the ALC). Importantly, *trans* complementation with p184INT_ALC in either ΔALC and H4 resulted in partial restoration of exoprotein but not the Yen-Tc (Fig. S8B,C) and cell morphology with cells of similar size to MH96 (Fig. S9). While *trans* complementation of ΔALC with either p184INT, p184 5’*holA*, and to a lesser extent with p184INT(K18) or p184ncALC220 resulted in cells of a similar length as observed for MH96 (Fig. S9). Further to this *trans* complementation with the intergenic region, in full or in part, including the ncRNA220, did not significantly alter cell morphology in H4 being similar in appearance to vector only control (Fig. S9). In K18 none of the *trans* complementation vectors altered either the observed exoprotein (Fig. 8D) or cell morphology (Fig. S9).

**Figure 8:**
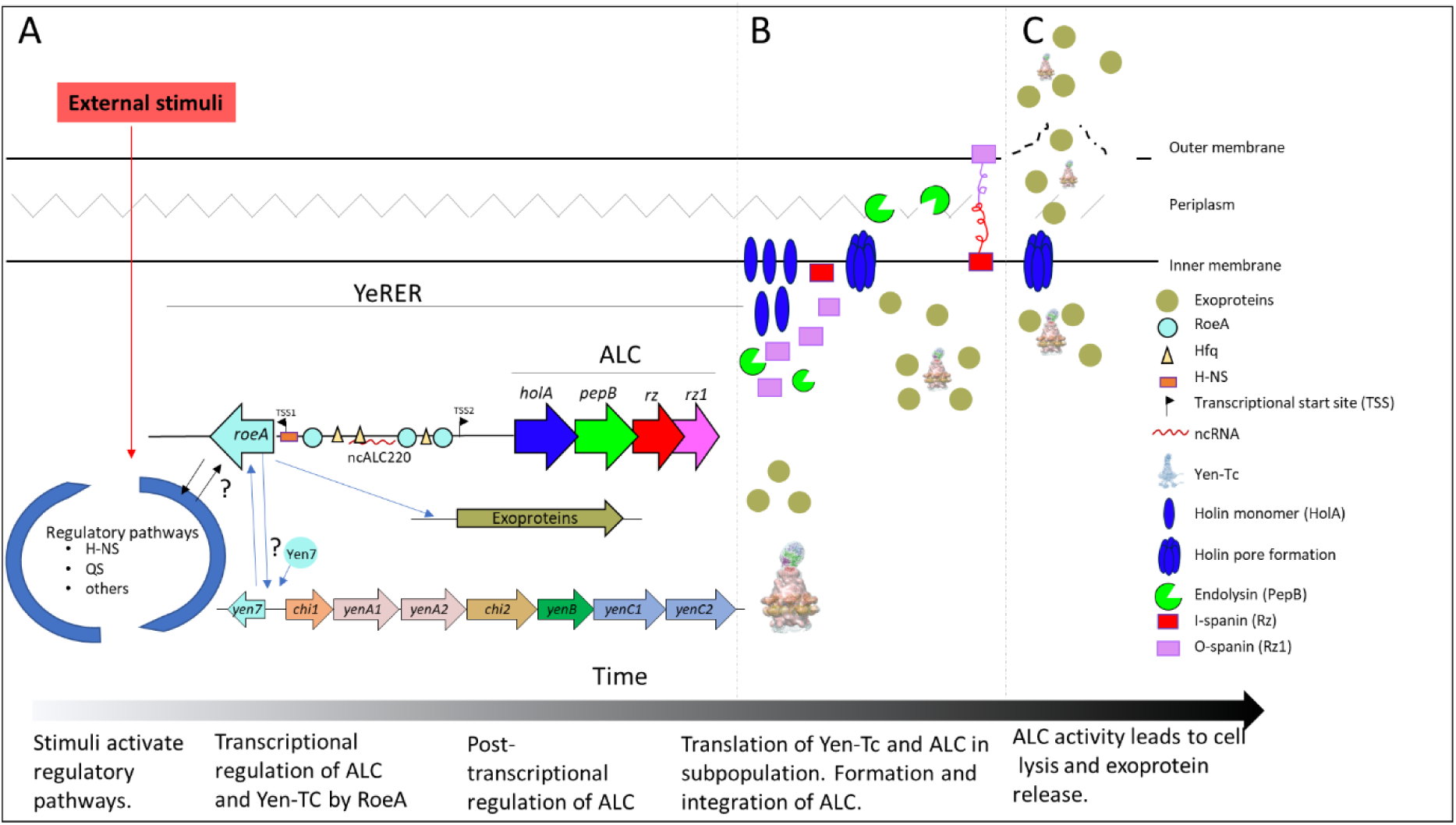
Schematic of the proposed model of *Y. entomophaga* exoprotein release. A) Devoid of a sensor domain, RoeA is likely activated as part of a larger regulatory system responding to environmental stimuli, such as temperature or stress response. The presence of a H-NS binding consensus sequence at similar nucleotide distances from *roe*A and its *yen*7 orthologue, it is likely H-NS regulates the expression of these genes. The RoeA-DNA binding motif allows its binding to PhoB-like DNA binding sites in the genome including the intergenic region of the YeRER. RoeA then activates the transcriptional expression of the ALC, and exoproteins including the Yen-Tc, as evidenced through their reduced transcription and the absence of pre-exoprotein in the in the H18 and H12 *roeA-* cell pellet. The ability of the ncALC220 region, but no truncation version thereof nor the intergenic region devoid of ncALC220, to reduce exoprotein release when placed *in trans* in MH96, suggests that the YeRER intergenic region may interact with mRNA or proteins e.g. Hfq, wherein Hfq binding sites reside within the YeRER intergenic region and within the ncALC220. B) The ALC is tightly regulated at post-transcriptional level by the termination-reinitiation complex of the ALC operon as well as yet to be determined post-transcriptional factors linked to ncALC220. Based on its observed expression in a subset of cells, HolA is under tight post-translational regulation. Prior to holin pore formation by a yet to be determined trigger, exoproteins accumulate in the cell causing cell elongation. Upon holin pore formation, the endolysin PepB and the spanin complex enter the periplasm, which subsequently C) causes cell lysis allowing the release of the exoproteins including the Yen-Tc. When no cell lysis occurs by inactivation of the ALC, such as in H4 and ΔALC, as evidenced by SDS-PAGE where pre exoproteins is observed in the cell pellet but not in the supernatant, causing cell elongation. In the H12 *roeA* mutant neither the ALC nor exoproteins are produced resulting in uniform sized cells which do not elongate.

To determine if cell elongation observed in H4 and ΔALC and cell shortening observed in K18 reflects an accumulation/absence of pre-exoprotein within the cell. The culture (∼ log_10_ 9.6 CFU/mL) cell pellets and supernatant of MH96, K18, H4, and ΔALC were assessed by SDS-PAGE (Fig. S10). Pre-exoproteins corresponding to the Yen-Tc (as proxy for exoproteins) were seen in the H4, ΔALC and MH96 but were absent in K18 and the *roeA* H12 mutant (Fig. S10A). To further define the role of RoeA in exoprotein release, the vector pAY-RoeA was induced in the *roeA* H12 and K18 mutants, where with the exception of the Yen-Tc, exoprotein was released (Fig. S10B).

## DISCUSSION

The presented data supports the role of the YeRER associated RoeA regulator and the ALC in production of exoprotein by *Y. entomophaga*. Differing to MH96, SDS-PAGE assessments of supernatant of the *roeA* mutants H12 and K18, and the ALC mutants ΔALC and H4 revealed an absence of exoprotein (Fig. S10) and changes in cell morphology (Fig. S9). The altered phenotype of reduced exoprotein release and changes in cell morphology in H4, ΔALC could be restored by *trans* complementation of p184INT_ALC, revealing that its putative RoeA regulator can act *in trans*. The presence of pre-exoprotein precursors such as the Yen-Tc in the cell pellet of ΔALC and H4 but no exoprotein (Fig. S10) supports the mechanistic role of the ALC in exoprotein release. Under the same conditions no pre-exoprotein was observed in the cell pellets of either the H12 *roeA* mutant or K18 (Fig. S10) further validating the regulatory role of RoeA in the induction of pre-exoprotein including the Yen-Tc.

The induction of the native ALC cassette pAY-ALCΔ*rz1* devoid of the second *rz1* spanin did not cause cell lysis. In *Y. enterocolitica* W22703 induction of its lysis cassette deficient in spanins caused cell lysis (37). However, as indicated in Fig. S1, W22703 *holA* and *pepB* are adjacent genes while in MH96 *holA* and *pepB* are overlapping genes on different reading frames. Through uncoupling the ALC system, the induction of pAY-ALC-opt, pAY-ALCΔ*rz-opt* (devoid of Rz spanin) and pAY-ALCΔ*rz1-opt* (devoid of the Rz1 spanin), caused rapid cell lysis in MH96, indicating the necessity of holin and endopeptidase for lysis. Similar effects were also noted on the induction of these constructs in *E. coli*. Of interest, although the induction pAY-ALCΔ*pepB-opt* devoid of *pepB* had no effect on MH96, its induction in *E. coli* resulted in a low level of cell lysis (Fig. 2C). This may reflect the unspecific pore of the holin allowing other peptidoglycan degrading enzymes to enter the periplasmic space in *E. coli* (38, 39).

Fluorescent microscopy of the HolA-sfGFP fusion found fluorescence was confined to a subset of cells of various sizes, wherein the fluorescence was observed throughout the entire cell. This may reflect the integration of HolA in the inner membrane and subsequent formation of the holin holomer required for release of the endopeptidase as documented in *E. coli* phage λ system and the mycobacteriophage D29 system expressed in *E. coli* prior to cell lysis (40, 41). In contrast, the florescence of the spanin Rz1-sfGFP was observed throughout the entire population and across the entire cell, which was also demonstrated assessing the spanin complex of the phage λ using a GFPΦRz fusion by Berry et al., (2012) (42). The post transcriptional regulation of ALC is likely through the need for back-translational coupling to uncouple the *holA* and *pepB* ORFs, the process of which was confined to a subset of cells as evidenced using the HolA-sfGFP reporter. Similar to phage systems such as the λ phage, the translation of HolA is likely a rate limiting step of cell lysis (43).

Through RNAseq the expression 2235 genes including those encoding components of the ALC cluster were significantly reduced in the H12 *roeA* mutant. This included the reduced transcription of several key exoproteins (PL78_18785, PL78_05495, PL78_11910, PL78_05310 (as validated through LC-ESI-MS/MS of wild type MH96 culture supernatant) and the Yen-Tc. In addition, through the use of P*_roeA_::lacZ* and the Yen-Tc P*_chi1_::lacZ* reporters, the β-galactosidase activities of both strains proportionally increased through exponential growth in LB broth at 25°C and were both significantly reduced at 37 °C linking the translation of both RoeA and the Yen-Tc.

Based on the presence of a PhoB-like DNA binding consensus sequence in RoeA, the altered expression of many genes of the H12 *roeA* mutant likely reflects the ability of RoeA to bind to multiple genomically encoded PhoB DNA binding sequences. Other genes significantly reduced in expression in the H12 *roeA* mutant included ribosomal genes, and genes involved in metabolism. Based on the absence of a ribosomal depletion step in our methodology (refer methods) these transcriptional changes may cause the cells to enter a state of reduced metabolic activity. The reduced metabolism in H12 and K18 may account for the observed homogenous, mid-exponential cell culture comprising cells of a shorter length relative to the heterogenous population with a small number of elongated cells observed in wildtype MH96 (Fig. S11). In contrast, a high number of atypically elongated cells were observed in the ΔALC mutant.

Interrogation of the *roeA/holA* intergenic nucleotide sequence revealed the presence of several nucleotide repeats, PhoB- and H-NS binding motifs, and secondary structures (Fig. S7B, C; Table S4). The prevalence of a range of different DNA binding motifs and the ncALC220 suggests that the YeRER intergenic region is the substrate for a complex of transcriptional, translational, and post-translational regulation (44) (Fig. S7B, Table S4). The positioning of a H-NS binding consensus sequence at similar nucleotide distances from *roe*A and its *yen*7 homologue (Fig. S7C, Table S4) suggests that RoeA is likely under control of the H-NS regulatory cascade, from where RoeA in turn acts on exoprotein and the ALC expression. This in part parallels the thermoregulation of the *Y. enterocolitica* W22703 LysR-like transcriptional regulator TcaR1/2 and H-NS (25, 26). Of interest, H-NS, a global transcriptional regulator, reacts to environmental cues (45, 46) and may indirectly take the place of a TCR response sensor that is absent in RoeA. Similar H-NS regulatory cascades have been described for H-NS in acid stress resistance in *E. coli* (45, 47). Adding to this, Schoof et al. (2022) (16) found transposon mutants H23 and H45 for quorum sensing (QS) N-acyl-homoserine lactone synthetase resulted in reduced exoprotein through the exponential growth phase (17), indicating growth-phase dependent regulation of RoeA resulting in exoprotein expressions.

A saliant finding was the ability of the intergenic ncALC220 region to reduce exoprotein release in MH96 when complemented. This reduction was not observed through complementation of the YeRER intergenic region devoid of the ncALC220 region (Fig. S8D). This data revealed the ncALC220 region as a key component of a complex regulatory network. Based on this it is tempting to speculate that the ncALC220 may inhibit transcription or translation of ALC or RoeA (48, 49). *Trans* complementation of p184INT, p1845’*holA*, p184INT(K18) and p184ncALC220 in ΔALC did not restore exoprotein release in ΔALC but did reduce the proportion of elongated cells, with cell sizes similar to those observed for MH96 (Fig. S9). Based on these findings, we hypothesise that the complemented regions are diluting out an activator of RoeA, therefore decreasing RoeA activity and therefore the production of intracellular pre-exoprotein (Fig. S10) and prevents cell elongation (Fig. S11). The inability of any complementation of intergenic region in full or in part, to alter the cell morphology (Fig. S9) or exoprotein profile of the K18 strain, revealed that the missing nucleotides of the K18 3 bp deletion, 131 bp 5’ of *roeA* are required for the expression of RoeA.

The absence of elongated cells in either the K18 or the roeA H12 mutants (Fig. S11) further supports the notion that RoeA is required to induce intracellular pre-exoprotein expression, which causes cell elongation. Of interest the induction pAY-RoeA restored exoprotein release to K18 and H12, where similar to the trans complementation of p184INT_ALC in ΔALC, no Yen- Tc associated bands were observed. We can only assume this may relate to a combination of factors, such as the a proximity of nc220 to its yet defined target, and/or most likely the regulation of the Yen7- RoeA homologue.

The heterogeneity of the observed cell shapes and HolA expression reflect observations in other systems. In *S. marcescens* and *E. coli* the expression of the exoproteins chitinase AB and colicin, respectively, caused cell elongation which was confined to a low proportion of cells (13, 50). In this respect 0.6%, 6% and 2% of *E. coli* cells that expressed colicin A, E2 and E7, respectively, were elongated (50). Using mKate fluorescence peptidase ChiX reporter, 1% of *S. marcescens* cells fluoresced which correlated to the co-expression of the ChiWXYZ lysis cluster and the ChiAB chitinases (13). The heterogeneous colicin expression is caused by bet-hedging, a risk spreading strategy in which stochastically occurring phenotypes of an isogenic population may adapt to changing environmental conditions (50, 51). Under *in vitro* conditions, exoprotein including Yen-Tc release is restricted to temperatures ≤25°C (35), which reflects a responsive switch rather than a stochastic switch (bet-hedging). In contrast to these *in vitro* findings, through the use of a YenA1 GFP reporter, elongated MH96 cells were observed *in vivo* in *G. mellonella* during early infection at 25°C but were absent at 37°C where only a limited number of cells fluoresced (18, 52), indicating an additional layer of complexity in the regulatory system.

Based on the results above, a model for the regulation of exoprotein production in *Y. entomophaga* is proposed in Figure 8.

Based on the phylogenetic data, RoeA-like regulators identified in some bacteria of the Enterobacteriaceae and Yersiniaceae may also enable the release of other large macromolecular toxin-transporting assemblies such as AFP/PVC complex of *P. chlororaphis*, through the activation of an ALC-like lysis cassette. Further research to define the role of RoeA and the ncRNA, ncALC220, in exoprotein regulation is required. The proposed model of *Y. entomophaga* MH96 mediated exoprotein release provides phenotypic evidence of the crucial role of a holin/endolysin based system in the programmed release of proteins. It provides further support to the roles of these lysis systems as T10SS, as initially proposed by Palmer et al (2021) using in silico approaches.

## MATERIALS AND METHODS

### Bacterial strains and culture conditions

Bacterial strains and plasmids used in this study are listed in Table S1 and Table S3, respectively. Bacteria were cultured in LB broth or on LB agar at 25°C (*Y. entomophaga*) or 37°C (*E. coli*). For *E. coli* ST18, the culture medium was supplemented with 50 µg mL^-1^ 5- aminolevulinic acid. Unless stated, cultures were incubated with shaking at 250 rpm in a Ratek model OM11 orbital incubator. Antibiotic concentrations used were (µg mL^-1^): *E. coli*, kanamycin (50), ampicillin (100); *Y. entomophaga*, kanamycin (100); spectinomycin (100), chloramphenicol (90), tetracycline (15).

### Molecular cloning

Standard DNA techniques were performed as described by Sambrook (53). Chromosomal DNA was isolated using PrepMan Ultra Sample Preparation Reagent (Thermo Fisher) and plasmid DNA was isolated using a High Pure Plasmid Isolation Kit (Roche). For amplification of genetic elements, Platinum *Taq* DNA polymerase (Invitrogen) was used according to manufacturer’s guidelines. Amplicons were purified using High Pure PCR Product Purification Kit (Roche). Primers are listed in Table S5.

When required PCR purified amplicons were ligated into pGEM-T Easy (Promega) following the manufacturer’s instructions. The resultant construct sequences were validated using M13_F and M13_R universal primers, or in other cloned constructs using construct specific primers (Table S5). Sequencing was performed using Macrogen Sequencing Services (Macrogen Inc., Seoul, Republic of Korea). DNA was electroporated into *Y. entomophaga* and its derivatives using the method of Dower et al. (54).

### Construction of ambiguous lysis cassette (ALC) mutant

For the deletion of the ALC, 2 kb 5’ and 2 kb 3’ of the ALC operon were PCR amplified using the primer pairs MS82/83 and MS84/85, respectively. The primers MS83 and MS84 harbor a complementary sequence to the pKD4 encoded kanamycin cassette that was amplified using the primers MS01 and MS02. The resultant amplicons were then assembled using fusion-PCR. The purified fusion-PCR amplicon ΔALC was cloned into pGEM to form pGEM-ΔALC, from where it was cloned into the suicide vector pJP5608 using SacI restriction sites (pJP5608- ΔALC). *E. coli* ST18 was used to conjugate pJP5608-ΔALC into MH96. Trans-conjugants were selected on LB-agar with kanamycin and tested for loss of pJP5608 tetracycline resistance. The final mutant MH96-ΔALC was subjected to PCR and sequence was validated using the peripheral validation primers MS29 and MS30.

### Construction of sfGFP reporter genes

Construction of sfGFP reporter genes. PCR amplicons 5’ HolA (MS88/MS89) and 3’ HolA (MS90/MS91) from MH96 and the amplicon sfGFP from pBAD::sfGFP (Table S3), were cloned together using fusion-PCR with primers MS88 and MS91. The fusion was then cloned into pJP5608 using XbaI and XmaI restriction sites to form pJP5806-HolAsfGFP. Fusion PCR of 5’ Rz1 (MS92/MS93), 3’ Rz1 (MS94/MS95) and sfGFP was used to amplify amplicon Rz1-sfGFP, which was then cloned into pGEMT-easy. To enable selection a spectinomycin cassette (MS96/MS97) was cloned into the Rz1-sfGFP BamHI restriction site and the final amplicon Rz1-sfGFP-Spec cloned into pJP5608, XbaI and XmaI restriction sites to form pJP5608- Rz1sfGFP. Trans-conjugants were selected on LB-agar for loss of pJP5608 tetracycline resistance and for Rz1-sfGFP growth on spectinomycin. The final strains were PCR and sequence validated using the peripheral validation primers MS98/MS99 and MS98/MS30 (Table S5).

### Construction of p184 *trans* complementation vectors

To construct the p184 vector series (Fig. 8) the various regions of the MH96 wild YeRER region were PCR amplified using PCR primers (Table S6). The resultant PCR amplicons were cloned into pGEM Teasy from where they were cloned into pACYC184 using the pGEMTeasy derived EcoRI restriction site. The K18 intergenic region was amplified (INT(K18)) from K18. Using GeneScript DNA of INT with a deletion of ncYLC220 (INTΔnvYLC220), and ncYLC220 truncations ncYLC80 and ncYLC110 flanked by EcoRV restriction were synthesized and used to clone into pACYC184 EcoRV to form the respective plasmids p184 INTΔnvYLC220, p184ncYLC80, and p184ncYLC110. The pACYC184 constructs were sequence validated with primer set MS/MS and transformed into MH96, K18, ΔYLC and H4.

### Protein visualisation and LC-ESI-MS/MS

Standard sodium dodecyl sulfate-polyacrylamide gel electrophoresis (SDS-PAGE) was performed as described (55). Proteins were visualized by silver staining according to Blum *et al.* (56).For electrospray ionization ion trap-tandem mass spectrometry (LC-ESI-MS/MS), the 0.1 % (w/v) SDS polyacrylamide gel was stained with Coomassie brilliant blue and the appropriate band was excised and prepared for LC-ESI-MS/MS spectrometry. Each gel band was analysed by mass spectrometry after de-staining, reduction with 0.1 M tris (2- carboxyethyl) phosphine (Fluka Chemie,GmbH, Buchs, Germany), alkylation with 20 μL of 0.15 M iodoacetamide (Sigma, St. Louis, MO, USA) and digestion for 18 hours with 1 µg of TPCK-trypsin (Promega, Madison, WI, USA) in presence of 10% acetonitrile (ACN). After digestion the peptides were dried and resuspended in 50 µl of 0.1% FA prior to injection on the mass spectrometer.

LC-ESI-MS/MS was performed on a nanoflow Ultimate 3000 UPLC (Dionex) coupled to maXis impact HD mass spectrometer equipped with a CaptiveSpray source (Bruker Daltonik, Bremen, Germany). For each sample, 1 µl of the sample was loaded on a C18 PepMap100 nano-Trap column (300 µm ID x 5 mm, 5- micron 100Å) at a flow rate of 3000 nl/min. The trap column was then switched in line with the analytical column ProntoSIL C18AQ (100 µm ID x 150 mm 3-micron 200Å). The reverse phase elution gradient was from 2% to 20% to 45% over 60 min, total 84 min at a flow rate of 600 nL/min. Solvent A was LCMS-grade water with 0.1% formic acid (FA); solvent B was LCMS-grade ACN with 0.1% FA.

The Q-TOF Impact HD (Bruker Daltonics) mass spectrometer was set up in a data-dependent automatic MS/MS mode where a full scan spectrum (50-2000 m/z, 2Hz) followed by 10 MS/MS (350 to 1500 m/z, 1-20Hz) of the most intense ions with charge states 2-3 selected.

### Genome sequencing of K18

Genomic DNA for genome sequencing was isolated using the ISOLATE II Genomic DNA Kit (Bioline). For identification of DNA alterations in strain K18, Illumina HiSeq 2500 System by Macrogen Sequencing Services was used. DNA sequences were trimmed using Trim_Galore (http://www.bioinformatics.babraham.ac.uk/projects/trim_galore/). Nucleotide differences were identified by alignment of Illumina reads against the MH96 genome sequence using the conda package for breseq version 0.33.0 with default parameters http://barricklab.org/twiki/bin/view/Lab/ToolsBacterialGenomeResequencing (57).

### Bioinformatic analysis

DNA sequences were trimmed and aligned against the genome of strain MH96 (GenBank accession number NZ_CP010029.1) using the Map to Reference function of Geneious Prime (58). Protein sequences were assessed using Phyre2 (http://www.sbg.bio.ic.ac.uk/phyre2/html/page.cgi?id=index) and BLASTP (https://blast.ncbi.nlm.nih.gov/Blast.cgi) with default settings. For gene synteny, the highest BLASTP hits to the query protein were used to pull genome data from the respective organism, covering 10 kb over the homologous genes. Multiple nucleotide alignments and Neighbour-Joining tree were then performed using standard settings in Geneious 10.0.9.

Amino acid alignments were performed using Geneious 10.0.9 using ClustalW and BLOSUM Matrix with a Gap open cost of 10 and a Gap extension cost of 0.1. Amino acid alignments were visualized using GeneDoc 2.7.000 (59).

### Cloning of ALC constructs and RoeA in pAY2-4 and their arabinose-based induction

Variations of the ALC operon were cloned into pAY2-4 *Nde*I and *Xho*I site. The PCR amplicon of *holA/pepB/rz* using primer pair MS101 and MS104 was cloned into pAY2-4 to form pAY-ALCΔ*rz1* (*hoLA, pepB, rz*). Optimised amplicons of the lysis cassette were synthesised at GeneArt (Thermo Fisher Scientific, USA) and designed to encode: i) *holA, pepB, rz,* and *rz1* as non-overlapping ORFs (pAY-ALC-opt), while maintaining ribosomal binding sites (*rbs)* and amino acid identity; ii) optimised regions encoding *holA, pepB, rz (*pAY-ALCΔ*rz1-*opt);iii) optimised region encoding *holA, rz, rz1* (pAY-ALCΔ*pepB-*opt); and *iv) holA,pepB,rz1* (pAY- ALCΔ*rz-*opt), refer to Table S2 for synthesised nucleotide sequence. Cells harbouring pAY-ALCΔ*rz1*, pAY-ALC- opt, pAY-ALCΔ*pepB-* opt, pAY-ALCΔ*rz-*opt and pAY-ALCΔ*rz1-*opt were grown in LB (40%) broth at 25°C and 200 rpm until an OD_600_ of 1 was reached. The cultures were induced with arabinose (0.2% final concentration), or the same volume of dH_2_O was added as control and placed at ambient temperature ∼ 22°C on a rotating platform at 40 rpm. The OD_600_ was measured every 15 min until 3 h and at 24 hpi.

For construction of pAY-RoeA the primers set MS65/MS66 were used to PCR amplify the amplicon RoeA, the purified product then cloned into pGEM T-easy and sequence validated with M13F/R primer. Using NdeI and XhoI cloning sites, the RoeA amplicon was cloned into the analogous sites of pAY-2 to form pAY-RoeA. Prospective pAY-RoeA clones were sequence validated using the AY_F and AY_R primers. pAY-RoeA was then electroporated into H12, K18 and. For pAY-RoeA induction, 50-mL cultures were grown in 40% LB to which 0.02% arabinose was added. The cultures were incubated for 16 h at 25°C under 250 rpm shaking from where samples were centrifugated at 8.000 × g for 5min to cellect cell pellets and culture supernatant to assess using SDS-PAGE.

### Light and fluorescent microscopy

For light microscopy, three µL of a MH96 cell culture at 16 hpi were observed under phase contrast. For Live/Dead staining the Syto9/PI stain (LIVE/DEAD *BacLight* kit; Invitrogen, Carlsbad, CA, USA) was used at a 1:1 ratio and incubated for 5 min in a 1.5 mL UV-safe tube. Cells were observed under an Olympus BX50 light microscope at × 400 magnification for both light and fluorescence microscopy. The SYTO9 stain was visualised using a FITC filter with excision of 460/515 nm, and the PI stain using Texas red 545/610 nm filter. Cell counts were measured using the software ImageJ 1.47v (60).

### Live-cell imaging

All cultures were grown in 100% LB broth (200 µg mL^-1^ ampicillin) until OD=1. For live-cell imaging, 10 µL of the culture were induced with 0.6% arabinose and immediately pipetted onto agarose covered glass slides and assessed by light-microscopy within the first minute post induction. Agarose pads were used to eliminate cell movement during the imaging process. Live-cell imaging was undertaken using the LSM710 microscope operated with Axiovision System (Carl Zeiss, Germany).

### Transmission electron microscopy (TEM)

Three µL of a 16-hpi culture sample was pipetted on carbon coated (3 nm) copper grids (200- mesh) (ProSciTech, Thuringowa, Australia) and incubated at ambient temperature (22°C) for 60 s. Residual liquid was removed with Whatman filter paper grade 1. For the negative stain, the sample was incubated with three µL of 0.7% uracil acetate for 45 s. After removing the residual liquid, the grids were dried at ambient temperature for at least 30 min and then examined in a Morgagni 268D transmission electron microscope (FEI, USA). Images were taken using an Olympus Megapixel III digital camera imaging system. Cell length and width was measured using the software ImageJ 1.47v.

### RNA isolation

RNA isolation was performed using the RNA Mini Kit (QIAGEN, Germany) and the corresponding RNAprotect Bacteria Reagent (QIAGEN) and RNase-Free DNase Set Kits (QIAGEN).

Three culture flasks per strain (H12, MH96) were incubated to reach log_10_ CFU mL^-1^ of 9.5. From each culture flask. 1 mL of sample culture was immediately transferred into 2 mL RNA protect bacteria (QIAGEN) and vortexed. After 5 min incubation at 25°C, the samples were pelleted at 5,000 × g for 10 min. The supernatant was decanted, and pellets left to air dry at 37°C before freezing at −20°C.

RNA was isolated using the RNA Mini Kit (QIAGEN) following the manufacturer’s instructions. Following the on-column DNA digest with RNAse-free DNAse, a second, off-column DNA digest was performed. To 40 µL RNA, RDD buffer (40 µL) and DNase stock I (2.5 µL) was added, and the volume adjusted to 100 µL with DNA-free water. After 10 min incubation time at 25°C, the RNA clean-up protocol (provided in the QIAGEN RNA Mini Kit) was followed. The RNA was eluted in 40 µL RNAse-free water and isopropanol precipitated. The total volume was adjusted to 180 µL and 1% sodium acetate (3 M) was added. Three times ice cold 100% ethanol (600 µL) were added to the solution and vortexed. The microcentrifuge tube was then placed at – 20°C overnight, after which the suspension was centrifuged (10,000 g for 30 min at 4°C) and supernatant discarded. The pellet was washed twice with ice cold 75% ethanol (500 µL) and pelleted at 10,000 × g for 5 min at 4°C and the supernatant discarded. After the final wash step the samples were pulse spun to remove residual ethanol by pipetting out residual supernatant. The pellets were air dried at 37°C for 30 min and then resuspended in RNase-free water. The resuspended sample was quantified by nanodrop. Of the sample RNA, 6 mg/µL were placed into a reaction tube, and liquid was evaporated in a SpeedVac. RNAseq was quality controlled and performed by Macrogen (South Korea).

### RNAseq

The Illumina short reads were inspected for quality using FASTQC (https://www.bioinformatics.babraham.ac.uk/projects/fastqc/). Bases with low quality PHRED scores (PHRED < 15 using a sliding window of 4 bases) were trimmed (using TRIMMOMATIC) from the short-read library, as well as any Illumina adapter sequences. Paired reads that were longer than 36 bp were kept for further analysis. The *Y. entomophaga* MH96 genome (Aug2018.NCBI.gb) was converted from GenBank format to fasta format using a custom BIOPYTHON script prior to indexing. Gene annotations were converted to gff format using the BIOPERL program bp_genbank2gff3.pl (https://manpages.debian.org/testing/bioperl/bp_genbank2gff3.1p.en.html). The genome fasta file was indexed and the short-read libraries aligned to the reference genome using HISAT2 with the default parameters. STRINGTIE was used for novel transcript assemblies, and BALLGOWN calculated the transcript count matrix for each sample.

The transcript count matrix was read into R and differential gene expression calculated using the DESeq2 package (61). Genes were considered differentially expressed when the adjusted p-value (p_adj_) was less than 0.05. For analysis purposes differently expressed genes of a log2Fold change <-1 and >1 were considered of interest and further assessed.

### β-galactosidase assay

For β-galactosidase (β-gal) assays an over-night culture was used to inoculate (1%) 50-mL LB flasks that were incubated at either 25 or 37 °C with 200 rpm shaking. For the assay, at each time point 2 x 200 µL of each culture were collected into a sterile 96-well plate (F-bottom) (Greiner Bio-One Cellstar) and then frozen at −80 °C. Using a 96-well microplate reader SPECTROstarNano (BMG Labtech) and the MARS Data Analysis software (BMG Labtech), the rate of β-galactosidase production was measured at OD_420_ following the methods of Schaefer et al. (62, 63) with a custom β-galactosidase mix: 60 mM Na_2_HPO_4_, 40 mM NaH_2_PO_4_, 10 mM KCl,1mM MgSO_4_; 1.8 μl mL^-1^ β-mercaptoethanol, 0.2 mg mL^-1^ Lysozyme from chicken egg (Sigma), 1:150 diluted Bacterial Protein Extraction Reagent (Thermo Fisher); and 1 mg mL^-1^ of 2-nitrophenyl-β-galactopyranoside (Sigma)). To control for temperature-dependent differences in the MH96 calibration curves measured at OD_600_ for 25 and 37 °C, an appropriate calibrating factor was applied to the Miller Unit Equivalent calculation.

## Data availability

Sequencing data was deposited in the NCBI databank with the BioProject PRJNA892653 with BioSample association numbers SAMN31393954, SAMN31393955, SAMN31393956 for MH96 transcriptome reads (MH1, MH2, MH3) and SAMN31393957, SAMN31393958, SAMN31393959 for H12 (*roeA* mutant) reads (RoeA1, RoeA2, RoeA3).

## ACKNOWLEDGMENTS

We thank Ruy Jauregu for bioinformatic Hiseq assessment of the K18 strain, Sandra Jones and Ancy Thomas for LC-ESI-MSMS of the MH96 supernatant. We also want to thank Martin Pilhofer and Miki Feldmüller from the Institute of Molecular Biology & Biophysics, Eidgenössische Technische Hochschule, Zürich for the providing tools and critical analysis of live-cell and fluorescent microscopy imaging. This work was supported by the Ministry of Business, Innovation and Employment, New Zealand (Next Generation Bio-pesticides, grant number C10X1310), Callaghan Innovation R&D fellowship grant and support from the Bio- Protection Research Centre.

