## Supplementary material for "Lysis cassette-mediated exoprotein release in *Yersinia entomophaga* is controlled by a PhoB- like regulator": Supplemetary figures and tables.pdf

Short title: *Regulation of* exoprotein release in *Y. entomophaga*

[#]Corresponding author:

Marion Schoof, AgResearch, Private Bag 4749, Christchurch 8140, New Zealand, +64 3 321 8740,

Mark RH Hurst, AgResearch, Private Bag 4749, Christchurch 8140, New Zealand, +64 3 325 9919,

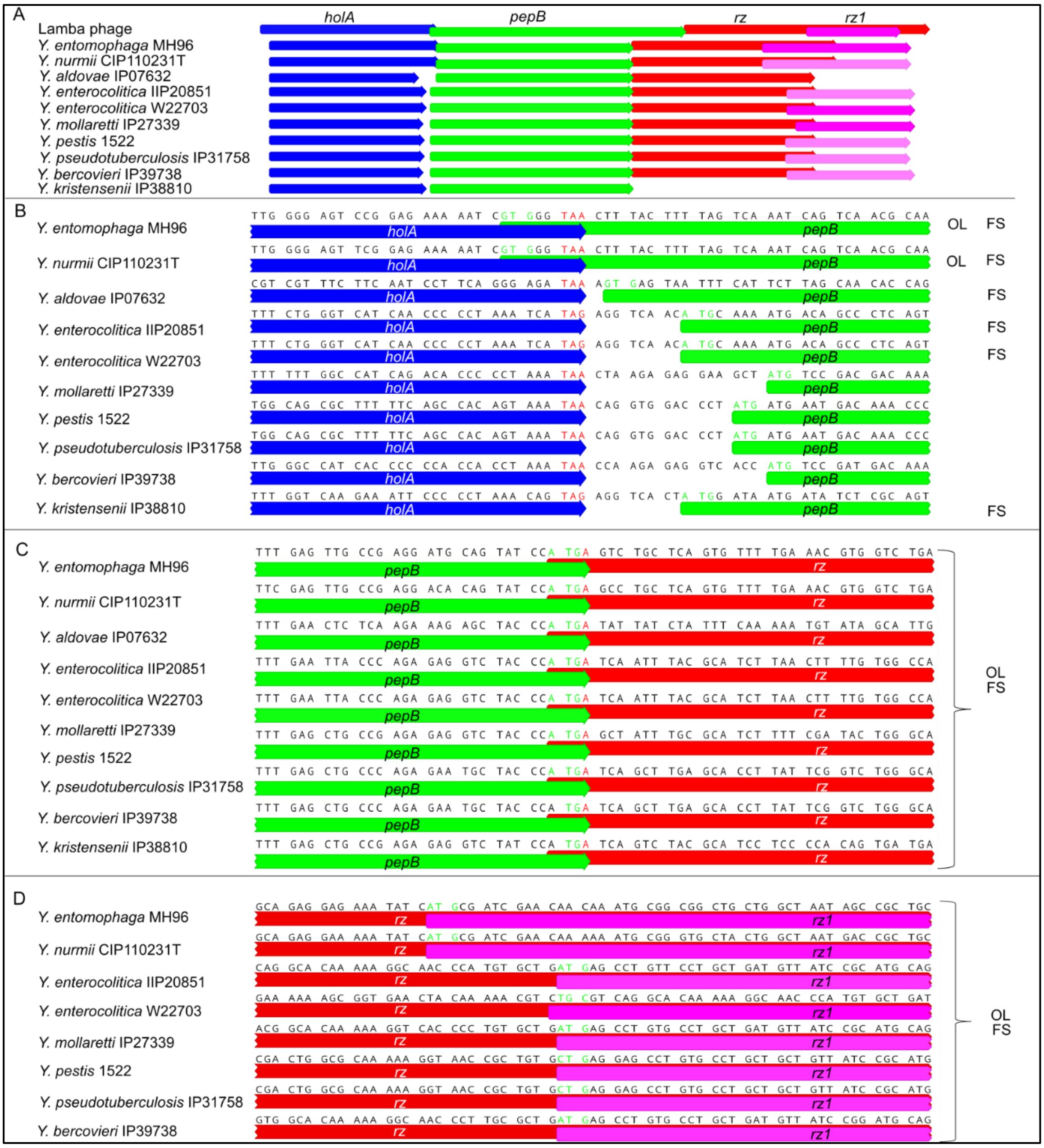


Figure S1: Arrangement of the ALC genes. A) Genetic arrangement of the lysis cassette genes after manual assessment of *rz1* annotation. Pink arrows denote the newly annotated Rz1 genes sharing >70% amino acid identity to the W22703 Rz1 gene.B-D) Nucleotide alignments showing overlap of the *holA, pepB* and *rz* of ALC-like lysis cassettes within different *Yersinia* species. Indicated are frameshifts (FS) and gene overlaps (OL) of the respective gene arrangement. B) Comparison of the *holA-pepB* gene arrangements. C) Conserved gene arrangement between *pepB-rz.* D) overlap *rz* and *rz1*.

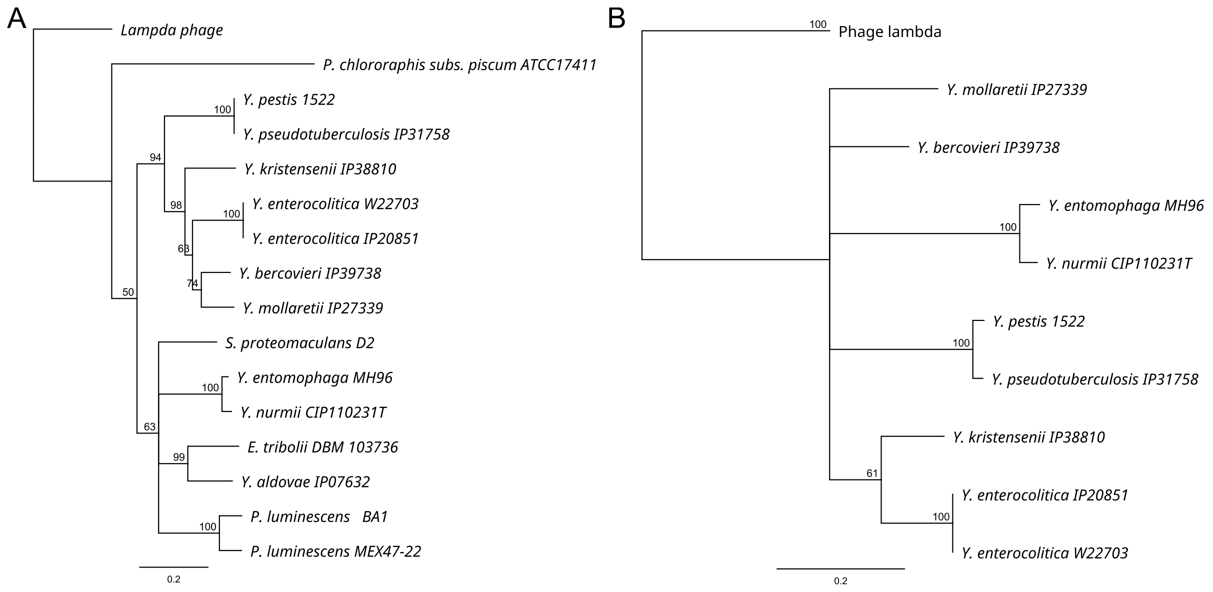


Figure S2: Phylogeny of the ALC encoded HolA (A) and (B) of PepB; maximum likelihood, 1000 bootstrap replicates, bootstrap values indicated at respective nodes, scale bar denoting patristic distances.

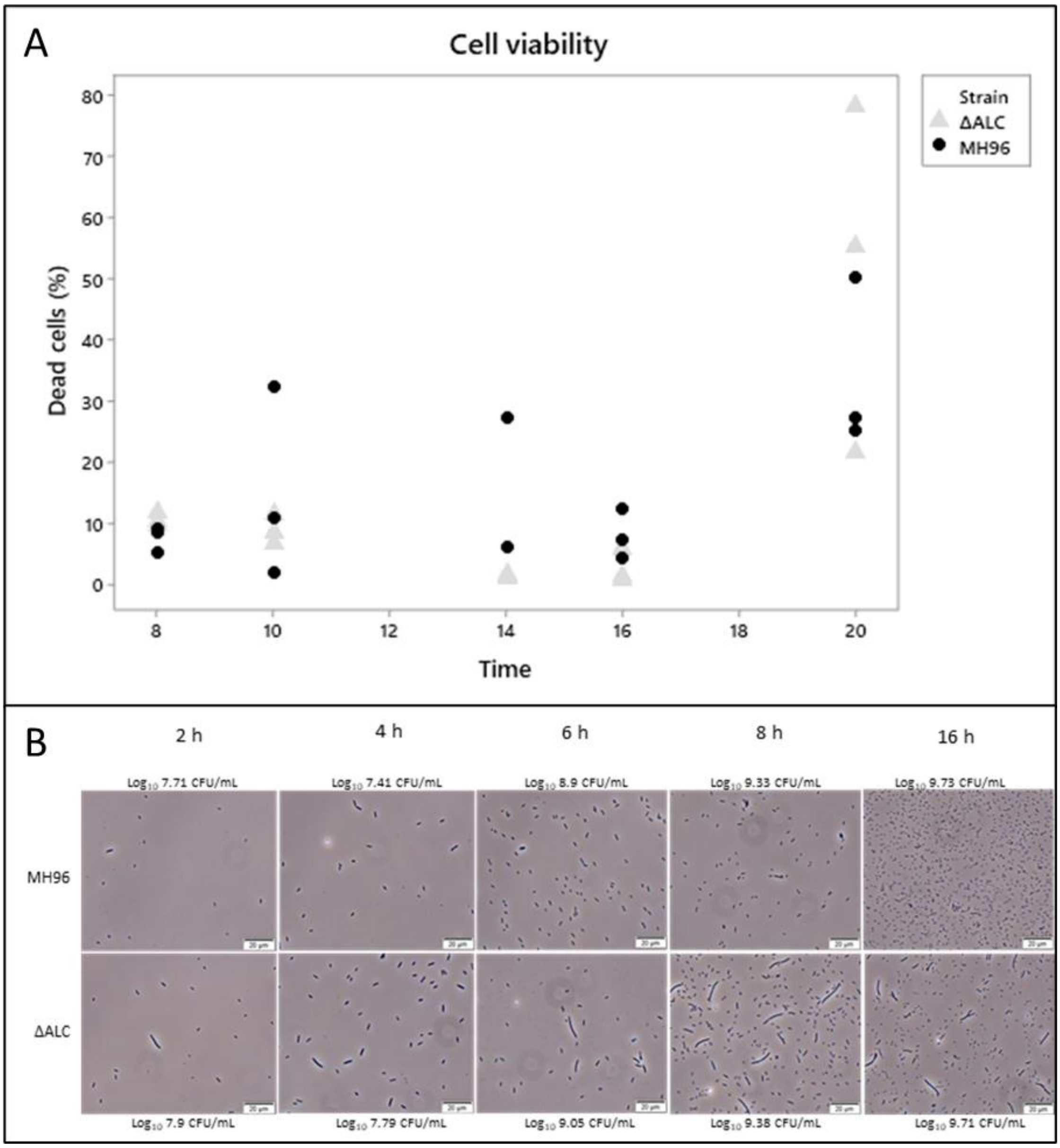


Figure S3: Cell viability of ΔALC. A) Cell viability assessed using the LIVE/DEAD stain from cultures taking at same time points as MH96. Cell viability is presented as proportion to the total cell count measured in 9 samples per strain at each time point. B) Light microscopy of time-dependent cell morphology and CFU of MH96 and ΔALC. The increasing proportion of elongated ΔALC cells can be seen over time. Scale bar (20 µm).

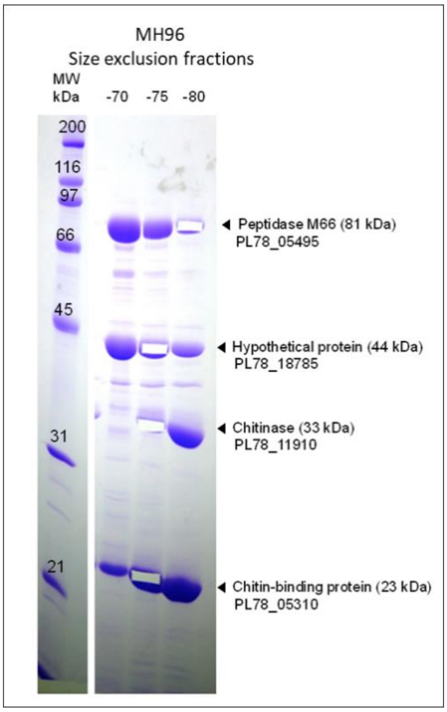


Band 1: Peptidase M66 (PL78_0545)

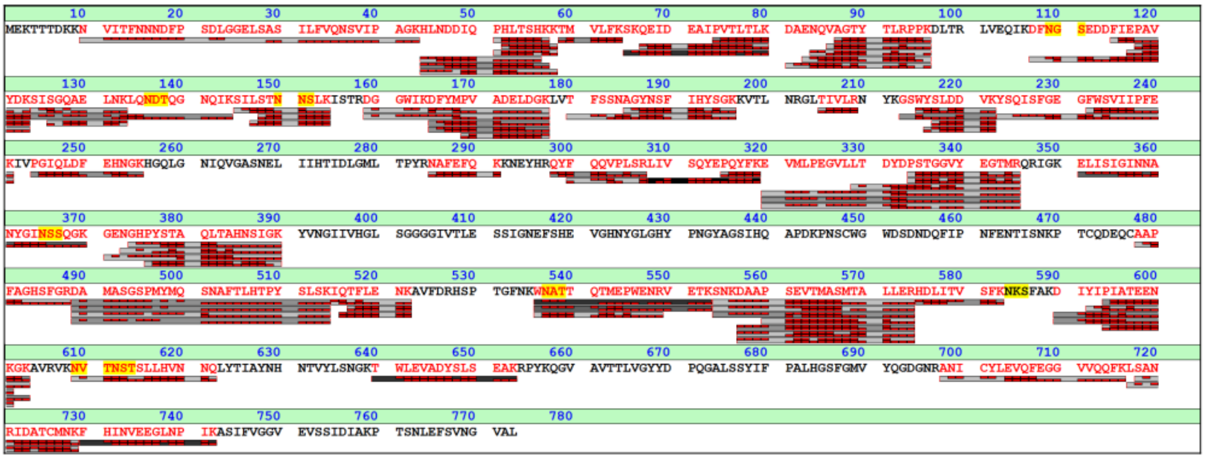


Band 2: Hypothetical protein (PL78_18785)

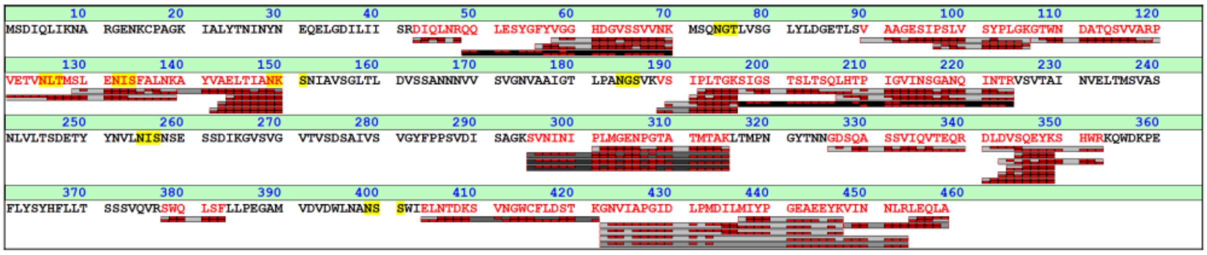


Band 3: Chitinase (PL78_11910)

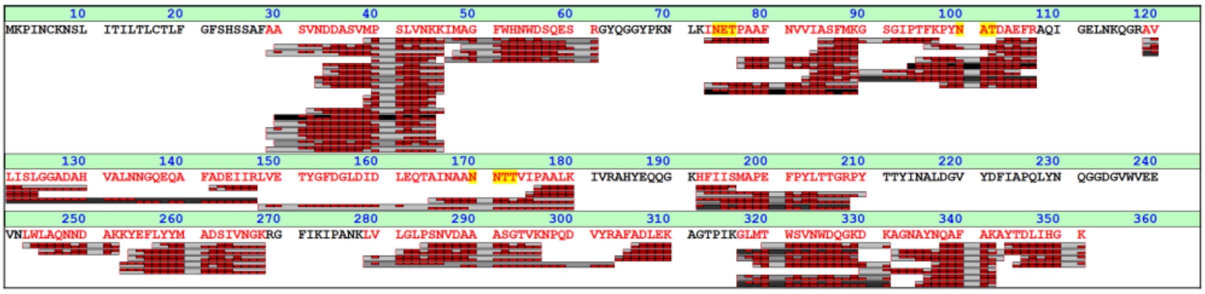

Band 4: Chitin binding protein (PL78_05310)

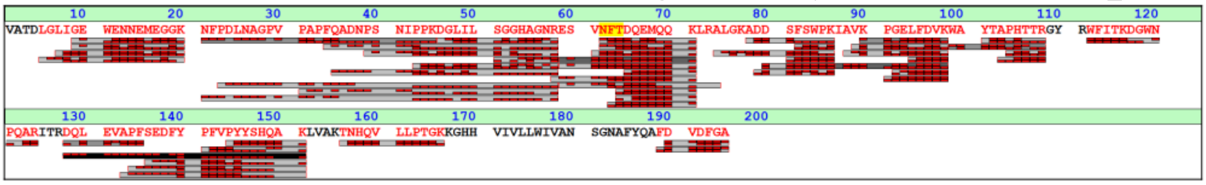


Figure S4: Size-exclusion chromatography fragment pools from cell supernatant of *Y. entomophaga* MH96 visualized on 12 % polyacrylamide gel by SDS-PAGE and stained with Coomassie brilliant blue. Excised bands are indicated by black arrows and the resultant LC-ESI-MS/MS data listed.

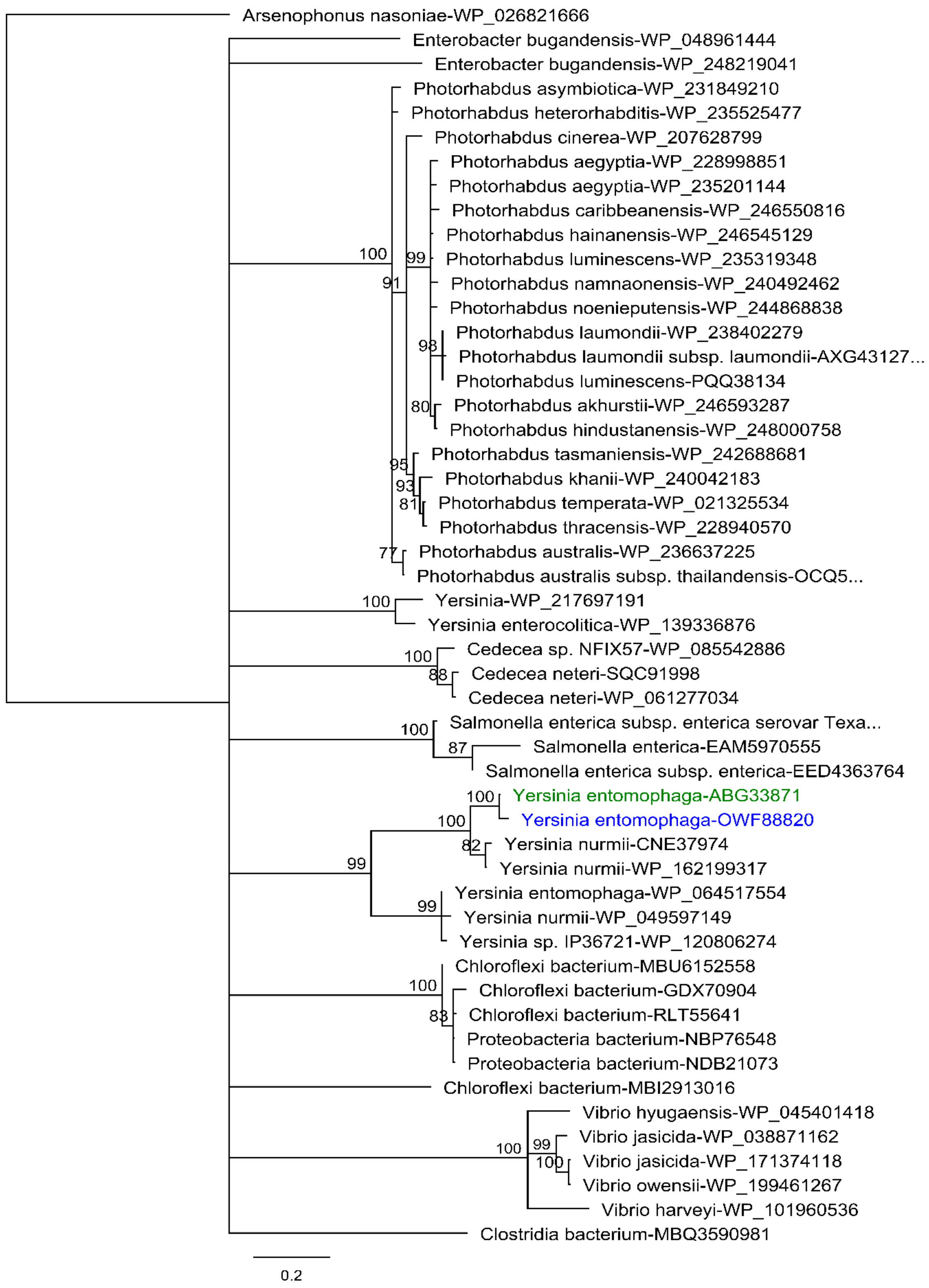


Figure S5: Phylogenetic analysis of RoeA orthologs. Protein sequences were aligned using MUSCLE and a phylogenetic tree was inferred using the neighbour-joining method, with 1000 bootstrap replicates indicated at respective nodes. Closest *Y. entomophaga* RoeA (blue) ortholog is *Y. entomophaga* Yen7 (green), which form a clade with *Y. nurmii*. Scale bar denoting patristic distances.

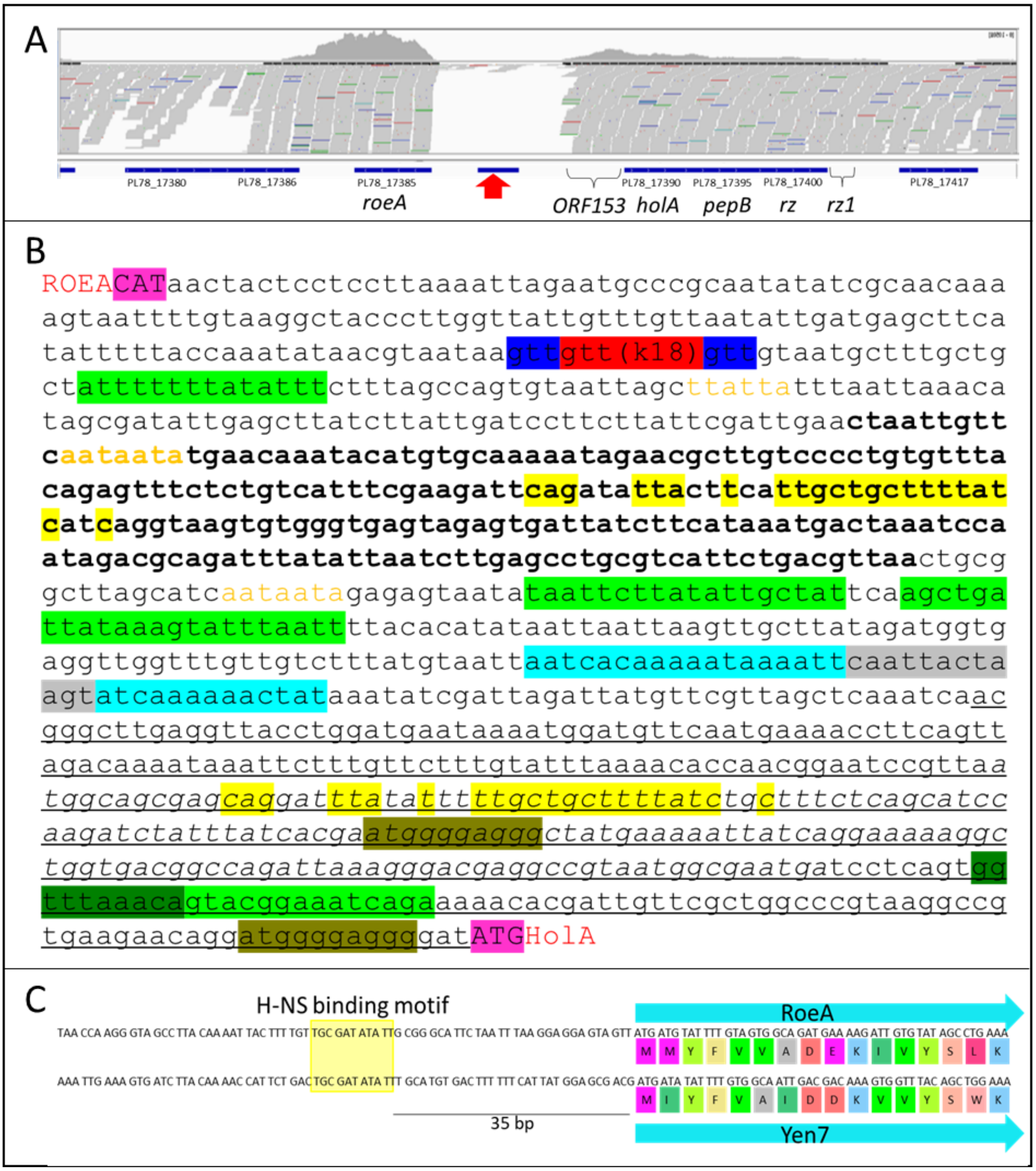


Figure S6: Analysis of YeRER intergenic region (INT). A) Read mapping of *in vitro* transcriptome analysis of the YeRER in MH96. Vertical red arrow denotes location of the predicted ncRNA (ncALC220); Location of predicted ORF153. B) Adjacent gene *holA* and *roeA* indicated (red letters). Hfq binding motifs based on Lorenz et al. 2010 (1) in orange letters. 5'UTR of *holA* indicated (underlined). Highlighted are the following: base pair deleted in K18 (red) within a 3 bp repeat region (dark blue); degenerate long repeat (yellow); hairpin structure (light blue) with loop (gray); PhoB and PhoB-like DNA binding motif (light green) – overlapping PhoB-binding motifs were identified within the hairpin structure and for visualization purpose not highlighted here; H-NS binding motif (dark green); perfect repeats (kaki). C) Homologous H-NS binding sequence 5' *roeA* and *yen7* based on Lang et al. (2007) (2).

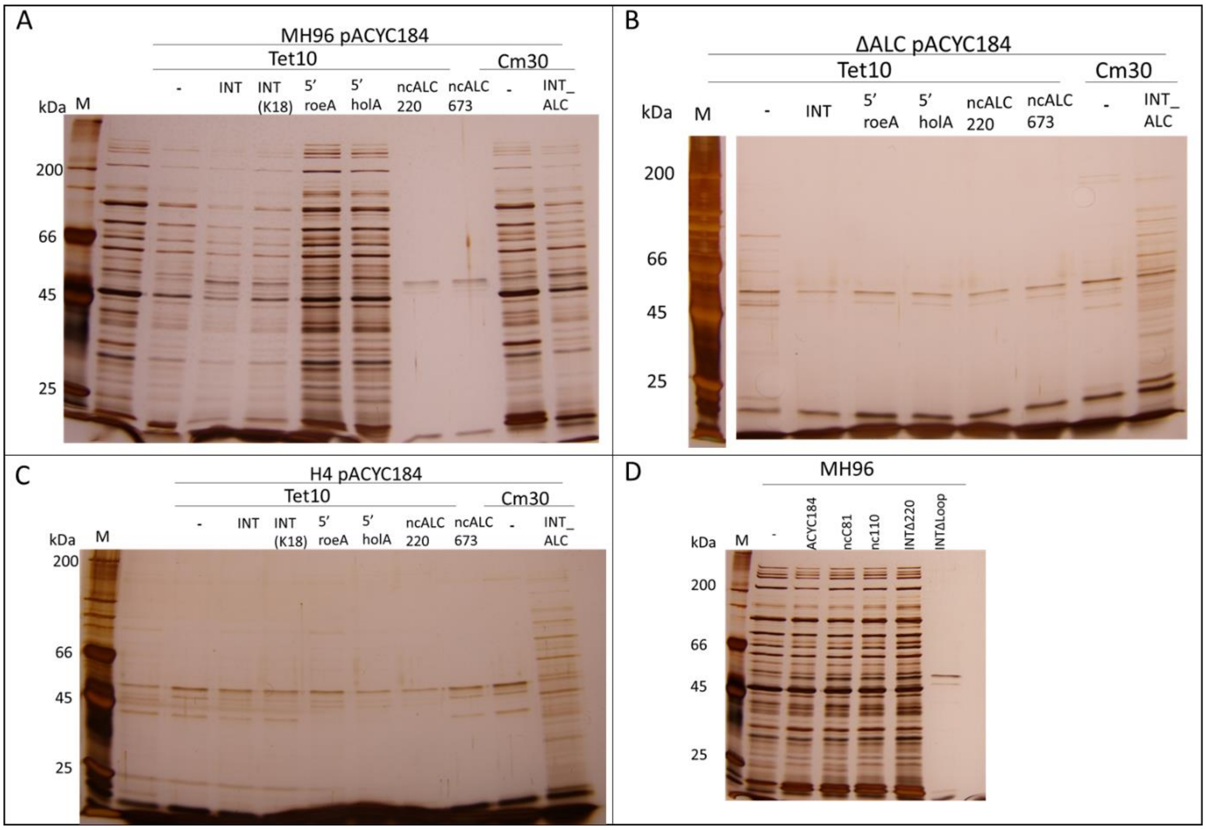


Figure S7: 10% SDS-PAGE of the *trans* complementation of p184 constructs in MH96 (A,D), ΔALC(B) and H4 (C). Samples are taken from supernatant of late exponential phase cell cultures. Antibiotic treatments for A,B,C are indicated. Tet indicates tetracycline 10 µg/mL and Cm indicates chloramphenicol 30 µg/mL. Cells of (D) were grown with chloramphenicol 30 µg/ml, except for wild type MH96 (-) which was grown without antibiotics.

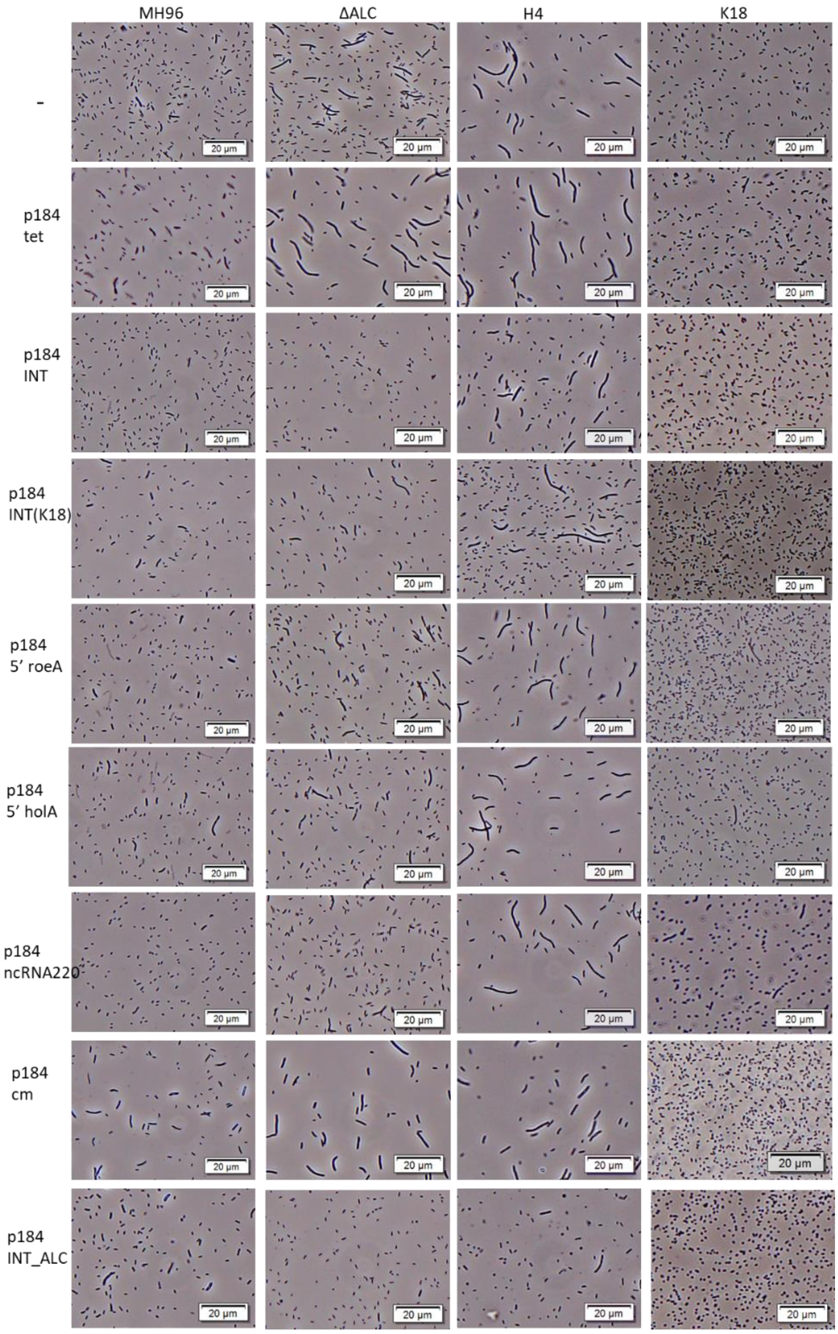


Figure S8: Light microscopy of in *trans* complementation with the listed p184 vector series (p184INT, p184 INT(K18), p184 5’*roeA,* p1845’*holA,* p184ncALC220, p184ncALC673, p184INT_ALC) used for *trans* complementation of the YeRER and the pACYC184 ( p184)control vector grown in the presence of 10 µg/mL tetracycline (tet) or 30 µg/mL chloramphenicol (cm) in either MH96, K18, H4 (5’ UTR *holA* mutant) and ΔALC. MH96, K18, H4 and ΔALC without vector are shown and indicated by (-). 20 µm scale bar indicated.

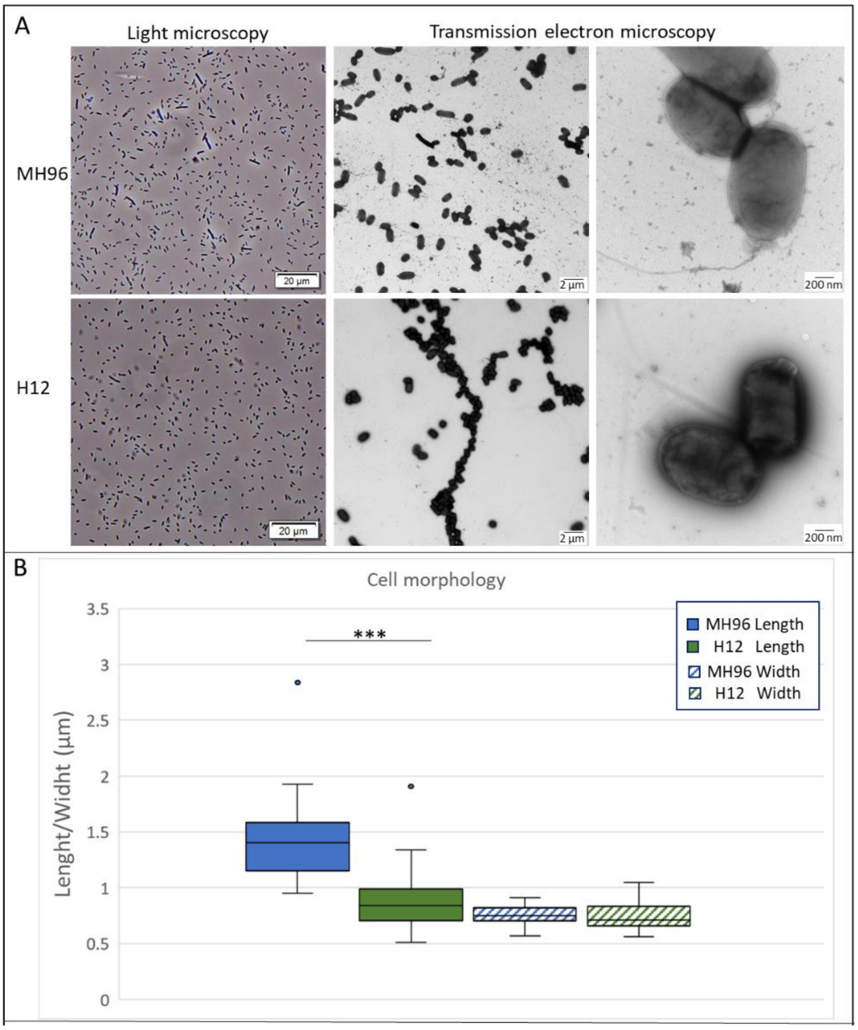


Figure S9: Cell morphology and exoprotein production of MH96 and the *roeA* mutant H12 in early stationary phase. *A)* Phase contrast- light microscopy and electron micrograph of MH96 and K18 cultures. Scale bars indicated. *B)* Cell size of MH96 and K18 measured in Length and Width. *** indicates p<0.0005, calculated by unpaired, two-sided t-test. C) Silver stained SDS-Page of cell pellet and supernatant of MH96, the YeRER transposon mutants H4 and H12 as well as ΔALC that are used in this study. Protein ladder (M) and size in kDa are indicated. Red boxes indicate absence of YenTc bands and red arrows indicate missing or reduced protein bands compared to MH96. Black arrows denote YenTc associated YenA bands in MH96, H4 and ΔALC cell pellets.

Table S 1: Bacteria strains and plasmids used in this study

| Strains | Description | Reference |
|---|---|---|
| *E. coli* | | |
| ST18 | *Escherichia coli* S17 λpir Δ*hemA* | (3) |
| DH10β | F- *mcrA* Δ(*mrr-hsdRMS-mcrBC*) Φ80d*lacZ*ΔM15 Δ*lacX*74 *endA1 recA1 deoR* Δ(*ara,leu*)7697 *araD*139 *galU galK nupG rpsL* λ- | (4) |
| CM2929 | $Cm^R$, *dam*13::Tn*9*, *dam*-6 | (5) |
| *Y. entomophaga* | | |
| MH96 | Wild-type strain, isolated from diseased *Costelytra giveni* larva | (6) |

| | | |
|---|---|---|
| K18 | Spontaneous MH96 non-secreting derivative, isolated at 4 weeks post field trial application. | AgResearch culture collection |
| H4 | Transposon mutant of MH96 generated by Tn5 insertion in 120 bp 5’ *holA* | (7) |
| H12 | Transposon mutant of MH96 generated by Tn5 insertion at 138 bp within *roeA* | (7) |
| MH96Δ*roeA151::Spec* | Sp, *roeA* mutation by gene disruption using sp-cassette insertion at position 151 nt of *roeA* | This study |

Table S2: GeneArt (GA) (Thermo Fisher Scientific, USA) and GenScript (GS) (USA) synthesized ALC constructs which were cloned into pAY. Underline indicates restriction sites, italics indicate ribosomal binding site and bold indicates start codon. Green highlights indicate HolA coding region, blue highlights indicate PepB coding region, kaki highlight indicates Rz coding region and yellow highlight indicates Rz1 coding region.

| | |
|---|---|
| ALC -opt<br><br>(GA) | CAT**ATG**ACGTTAAAGATTCCGACTGATACACAGGCAATTACCTGGTTAATTATTGGCCTTTTTTCAG<br>CTTGGGGAGGTGTTGTGAGATATCTCATGGATATGCAAGGGCACAGGGGGAGTTGGAGTTGGAT<br>GGGAATAATAAGTCAGATCATTATTTCAAGCTTCACTGGCCTTTTGGGGGGATTACTCAGCTTTGA<br>AAATGGCAGTAGTCATTACATGACTTTTGCTATTGCAGGCTTATTTGGCACCTTGGGGAGTACGGC<br>GTTGAGTTACCTGTGGCGGCGTTTCTTGGGGAGTCCCGAAAAGAACCGTGGGTAAAG*GAGGA*AA<br>AAC**GTG**GGTAACTTTACTTTTAGTCAAATCAGTCAACGCAATCTACAAGGCATACACCCGGATCTC<br>GTTCGGGTGGTGTACCTGGCACTCAAATTGTCCGAGGTGGATTTTCGCGTTATCGAAGGGCTGCG<br>TGATAGCGCACGTCAACGGCAAATGGTGCTTAATGGCAAGAGCCAAACGCTCAATAGCCGGCATC<br>TTACCGGTCATGCGGTCGATTTGGCGCCTTGGTTCAACAACACGATTCCCTGGAACGATTGGGGA<br>GCGTTTGCTCAGGTTGCAGCGGCGATGAAACAGGCGGCGAAGCAACTACAAATTCCTGTGATCTG<br>GGGCGGTGACTGGACGACGCTTAAGGATGGGCCGCACTTTGAGTTGCCCCGTATGCAGTATCCTT<br>GAA*GGAGGA*AAAAA**ATG**AGTCTGCTCAGTGTTTTGAAACGTGGTCTGCTACTTGCCACGCTATTT<br>GCGGTTTTGGTTGGCGGGCTGTGGGTTGCTCAACTGAAAAAGACCGCGACATTACTGAATTCTGA<br>AAATATCAGTCTGACGCAGCAGAATCAGCTTTATCTGCAACGGTTGCAAGGATATGACCAGCAGA<br>TGAAAACACTGGATGAGGCGTTAACCACTAATGCCATAACTCAGCGTCAGGCAGAGGAGAAATAT<br>CATGCGATCGAACAACAAATGCGGCGGCTGCTGGCTAATAGCCGCTGCGCTCGGGAGCCTGTGC<br>CTGATGCTGTTATCCGGTTGCAGCAACGAGCACTTGGCCAAAGCCTGCCCGGTGCAGCCAATAGT<br>ACCGAAAGCGCTGCTATTGCCCCATGACA*GAGGA*GAAATATC**ATG**CGATCGAACAACAAATGCG<br>GCGGCTGCTGGCTAATAGCCGCTGCGCTCGGGAGCCTGTGCCTGATGCTGTTATCCGGTTGCAGC<br>AACGAGCACTTGGCCAAAGCCTGCCCGGTGCAGCCAATAGTACCGAAAGCGCTGCTATTGCCCCA<br>TGATCCCCCGTTATTTAACGTCACAAAGTGGGGGGACTATCCCGGTTATGTGGAAGAATTAAATCT<br>ATCGTTGGCGCGTTGTAATGCAGATAAAACGGCGGTGGCCCAAATAATGTCTCCCCAAAAAGGAT<br>AATCCCGGATCCCTCGAG |
| ALCΔ*pepB*-opt<br><br>(GS) | CAT**ATG**ACGTTAAAGATTCCGACTGATACACAGGCAATTACCTGGTTAATTATTGGCCTTTTTTCAG<br>CTTGGGGAGGTGTTGTGAGATATCTCATGGATATGCAAGGGCACAGGGGGAGTTGGAGTTGGAT<br>GGGAATAATAAGTCAGATCATTATTTCAAGCTTCACTGGCCTTTTGGGGGGATTACTCAGCTTTGA<br>AAATGGCAGTAGTCATTACATGACTTTTGCTATTGCAGGCTTATTTGGCACCTTGGGGAGTACGGC<br>GTTGAGTTACCTGTGGCGGCGTTTCTTGGGGAGTCCCGAAAAGAACCGTGGGTAAA*GGAGGA*AA<br>AAA**ATG**AGTCTGCTCAGTGTTTTGAAACGTGGTCTGCTACTTGCCACGCTATTTGCGGTTTTGGTT<br>GGCGGGCTGTGGGTTGCTCAACTGAAAAAGACCGCGACATTACTGAATTCTGAAAATATCAGTCT<br>GACGCAGCAGAATCAGCTTTATCTGCAACGGTTGCAAGGATATGACCAGCAGATGAAAACACTGG<br>ATGAGGCGTTAACCACTAATGCCATAACTCAGCGTCAGGCAGAGGAGAAATATCATGCGATCGAA<br>CAACAAATGCGGCGGCTGCTGGCTAATAGCCGCTGCGCTCGGGAGCCTGTGCCTGATGCTGTTAT<br>CCGGTTGCAGCAACGAGCACTTGGCCAAAGCCTGCCCGGTGCAGCCAATAGTACCGAAAGCGCT<br>GCTATTGCCCCATGACA*GAGGA*GAAATATC**ATG**CGATCGAACAACAAATGCGGCGGCTGCTGGCT<br>AATAGCCGCTGCGCTCGGGAGCCTGTGCCTGATGCTGTTATCCGGTTGCAGCAACGAGCACTTGG<br>CCAAAGCCTGCCCGGTGCAGCCAATAGTACCGAAAGCGCTGCTATTGCCCCATGATCCCCCGTTAT<br>TTAACGTCACAAAGTGGGGGGACTATCCCGGTTATGTGGAAGAATTAAATCTATCGTTGGCGCGT<br>TGTAATGCAGATAAAACGGCGGTGGCCCAAATAATGTCTCCCCAAAAAGGATAATCCCGGATCCC<br>TCGAG |

| | |
|---|---|
| ALCΔ*rz*-opt (GS) | CAT**ATG**ACGTTAAAGATTCCGACTGATACACAGGCAATTACCTGGTTAATTATTGGCCTTTTTTCAG<br>CTTGGGGAGGTGTTGTGAGATATCTCATGGATATGCAAGGGCACAGGGGGAGTTGGAGTTGGAT<br>GGGAATAATAAGTCAGATCATTATTTCAAGCTTCACTGGCCTTTTGGGGGGATTACTCAGCTTTGA<br>AAATGGCAGTAGTCATTACATGACTTTTGCTATTGCAGGCTTATTTGGCACCTTGGGGAGTACGGC<br>GTTGAGTTACCTGTGGCGGCGTTTCTTGGGGAGTCCCGAAAAGAACCGTGGGTAAAG*GAGGA*AA<br>AAC**GTG**GGTAACTTTACTTTTAGTCAAATCAGTCAACGCAATCTACAAGGCATACACCCGGATCTC<br>GTTCGGGTGGTGTACCTGGCACTCAAATTGTCCGAGGTGGATTTTCGCGTTATCGAAGGGCTGCG<br>TGATAGCGCACGTCAACGGCAAATGGTGCTTAATGGCAAGAGCCAAACGCTCAATAGCCGGCATC<br>TTACCGGTCATGCGGTCGATTTGGCGCCTTGGTTCAACAACACGATTCCCTGGAACGATTGGGGA<br>GCGTTTGCTCAGGTTGCAGCGGCGATGAAACAGGCGGCGAAGCAACTACAAATTCCTGTGATCTG<br>GGGCGGTGACTGGACGACGCTTAAGGATGGGCCGCACTTTGAGTTGCCCCGTATGCAGTATCCTT<br>GACA*GAGGAG*AAATATC**ATG**CGATCGAACAACAAATGCGGCGGCTGCTGGCTAATAGCCGCTGC<br>GCTCGGGAGCCTGTGCCTGATGCTGTTATCCGGTTGCAGCAACGAGCACTTGGCCAAAGCCTGCC<br>CGGTGCAGCCAATAGTACCGAAAGCGCTGCTATTGCCCCATGATCCCCCGTTATTTAACGTCACAA<br>AGTGGGGGGACTATCCCGGTTATGTGGAAGAATTAAATCTATCGTTGGCGCGTTGTAATGCAGAT<br>AAAACGGCGGTGGCCCAAATAATGTCTCCCCAAAAAGGATAATCCCGGATCCGGATCCCTCGAG |
| ALZΔ*rz1*-opt<br>(GA) | CAT**ATG**ACGTTAAAGATTCCGACTGATACACAGGCAATTACCTGGTTAATTATTGGCCTTTTTTCAG<br>CTTGGGGAGGTGTTGTGAGATATCTCATGGATATGCAAGGGCACAGGGGGAGTTGGAGTTGGAT<br>GGGAATAATAAGTCAGATCATTATTTCAAGCTTCACTGGCCTTTTGGGGGGATTACTCAGCTTTGA<br>AAATGGCAGTAGTCATTACATGACTTTTGCTATTGCAGGCTTATTTGGCACCTTGGGGAGTACGGC<br>GTTGAGTTACCTGTGGCGGCGTTTCTTGGGGAGTCCCGAAAAGAACCGTGGGTAAAG*GAGGA*AA<br>AAC**GTG**GGTAACTTTACTTTTAGTCAAATCAGTCAACGCAATCTACAAGGCATACACCCGGATCTC<br>GTTCGGGTGGTGTACCTGGCACTCAAATTGTCCGAGGTGGATTTTCGCGTTATCGAAGGGCTGCG<br>TGATAGCGCACGTCAACGGCAAATGGTGCTTAATGGCAAGAGCCAAACGCTCAATAGCCGGCATC<br>TTACCGGTCATGCGGTCGATTTGGCGCCTTGGTTCAACAACACGATTCCCTGGAACGATTGGGGA<br>GCGTTTGCTCAGGTTGCAGCGGCGATGAAACAGGCGGCGAAGCAACTACAAATTCCTGTGATCTG<br>GGGCGGTGACTGGACGACGCTTAAGGATGGGCCGCACTTTGAGTTGCCCCGTATGCAGTATCCTT<br>GAA*GGAGGA*AAAAA**ATG**AGTCTGCTCAGTGTTTTGAAACGTGGTCTGCTACTTGCCACGCTATTT<br>GCGGTTTTGGTTGGCGGGCTGTGGGTTGCTCAACTGAAAAAGACCGCGACATTACTGAATTCTGA<br>AAATATCAGTCTGACGCAGCAGAATCAGCTTTATCTGCAACGGTTGCAAGGATATGACCAGCAGA<br>TGAAAACACTGGATGAGGCGTTAACCACTAATGCCATAACTCAGCGTCAGGCAGAGGAGAAATAT<br>CATGCGATCGAACAACAAATGCGGCGGCTGCTGGCTAATAGCCGCTGCGCTCGGGAGCCTGTGC<br>CTGATGCTGTTATCCGGTTGCAGCAACGAGCACTTGGCCAAAGCCTGCCCGGTGCAGCCAATAGT<br>ACCGAAAGCGCTGCTATTGCCCCATGA |

Table S3: Plasmids used throughout this study

| Plasmid | Description | Source or reference |
|---|---|---|
| pKD4 | Kan, cloning vector | (8) |
| pGEM T-Easy | Amp, cloning vector, LacZ multi-cloning site | Promega Ltd. |
| pJP5608 | Tet, suicide vector, | (9) |
| pAY2-4 | Amp, arabinose induction vector | (10) |
| pACYC184 | Cm, Tet, cloning vector | (11) |
| pHP45 | Spec, cloning vector | (12) |
| pBAD::sfGFP | Amp, sfGFP cloning vector | (13) |
| pGEMΔALC | Kan, Amp, cloning vector of deletion *holA/pepB/rz* by replacing with kanamycin cassette; ALC flanking regions 1.5 kb (ΔALC amplicon) | This study |
| pJP5608ΔALC | Tet, Kan, suicide vector, ΔALC amplicon | This study |
| pAY-ALC-opt | pARA, expression vector of optimised ALC coding region expressing *holA/pepB/rz/rz1* | This study |
| pAY-ALCΔ*rz1* | pARA, expression vector of *holA/pepB/rz* | This study |
| pAY- ALCΔ*rz1-opt* | pARA expression vector of optimsed ALC expressing *holA/pepB/rz* | This study |
| pAY- ALCΔ*rz*-opt | pARA expression vector of optimsed ALC expressing *holA/pepB/rz1* | This study |
| pAY- ALCΔ*pepB*-opt | pARA expression vector of optimsed ALC expressing *holA/rz/rz1* | This study |
| P184-INT | pACYC184, tet, expressing YeRER intergenic region from MH96 (1038 bp) | This study |
| P184-INT(K18) | pACYC184, tet, expressing YeRER intergenic region from K18 (1035 bp) | This study |
| P184-INT_ALC | pACYC184, cm, expressing YeRER intergenic region and *holA/pepB/rz* from MH96 (2133 bp) | This study |
| P184-5’holA | pACYC184, tet, expressing YeRER intergenic region of 5’ *holA* from MH96 (571 bp) | This study |
| P184-5’roeA | pACYC184, tet, expressing YeRER intergenic region of 5’ *roeA* from MH96 (467 bp) | This study |
| P184-ncALC220 | pACYC184, tet, expressing ncRNA from YeRER intergenic region (220 bp) | This study |
| P184-ncALC81 | pACYC184, tet, expressing truncated ncRNA from YeRER intergenic region (81 bp) | This study |
| P184-ncALC673 | pACYC184, tet, expressing partial YeRER intergenic region including ncRNA220 from MH96 (673 bp) | This study |

Table S4: Nucleotide protein binding sites located in the YeRER intergenic region of *roeA* and *holA* identified using CollecTF (14) and Prodoric programmes (15). Indicating binding sequence of known DNA-binding proteins in different gram-negative bacteria species, and location within the YeRER intergenic region.

| Binding protein | Position within intergenic ReYER (nt) | Binding sequence | Source strain(s) |
|---|---|---|---|
| Binding sequences in intergenic region of YeRER | | | |
| PhoP | 67 - 89<br>67 - 89<br>70-87 | CCTTGGTTATTGTTTGTTAATAT<br>CCTTGGTTATTGTTTGTTAATAT<br>TGGTTATTGTTTGTTAAT | *E. coli K12*<br>*Salmonella …*<br>*Y. pestis.* |
| Fur | 70-90<br>75-92 | TGGTTATTGTTTGTTAAT<br>TGGTTATTGTTTGTTAATATT | *Dyckia*<br>*Y. enterocolitica* |
| OmpR | 148-164<br>151-166 | CTGCTATTTTTTTATAT<br>CTATTTTTTTATATTT | *Y. enterocolitica 8081*<br>*Y. enterocolitica YPIII* |
| RegA | 153-170 | CTATTTTTTTATATTT | *Citrobacter* |
| ArcA | 155-175 | ATTTTTTTATATTTCTTT | *E. coli K12* |
| RdgB | 163-186 | ATTTCTTTAGCCAGTGTAATTAGC | *Pectobacterium carotovorum subsp. carotovorum PCC21* |
| OxyR | 155-188 | TTTTTTATATTTCTTTAGCCAGTGTAATTAGCTT | *Pseudomonas putida KT2440* |
| Fur | 485-505<br>488-518 | AATAATAGAGAGTAATATAAT<br>AATAGAGAGTAATATAATTCTTATATTGCTA | *Dyckia*<br>*E. coli K12* |
| RegA | 502-519 | TAATTCTTATATTGCTAT | *Cirtrobacter* |
| PhoP | 523-545<br>526-543 | AGCTGATTATAAAGTATTTAATT<br>TGATTATAAAGTATTTAA | *E. coli K12*<br>*Y. pestis* |
| PhoB | 523-545 | AGCTGATTATAAAGTATTTAATT | *Salmonella* |
| ArcA | 544-564 | TTTTACACATATAATTAATTA | *E. coli K12* |
| CsgD | 542-555 | AATTTTACACATAT | *E. coli K12* |
| ArcA | 591-611 | TTGTTGTCTTTATGTAATTAA | *E. coli K12* |
| Fur | 613-631<br>616-636<br>642-659 | CACAAAAATAAAATTCAAT<br>AAAAATAAAATTCAATTACTA<br>CAAAAAACTATAAATATC | *Y. pestis*<br>*Dickya*<br>*salmonella* |
| ToxR | 620-629 | ATAAAATTCA | *V. cholerae* |
| PhoP | 638-660<br>640-657<br>641-657 | GTATCAAAAAACTATAAATATCG<br>ATCAAAAAACTATAAATA<br>TCAAAAAACTATAAATA | *Salmonella*<br>*Y. pestis*<br>*E. coli K12* |
| MntR | 650-675 | TATAAATATCGATTAGATTATGTTCG | *E. coli K12* |
| Fur | 940-958 | ATGGCGAATGATCCTCAGT | *Y. pestis* |
| ArcA | 957-972 | GTGGTTTAAACAGTAC | *V. fischeri* |
| HNS | 959-968 | GGTTTAAACA | *E. coli K12* |
| PhoP | 960-982<br>960-982<br>962-980<br>962-980 | GTTTAAACAGTACGGAAATCAGA<br>GTTTAAACAGTACGGAAATCAGA<br>TTAAACAGTACGGAAATCA<br>TTAAACAGTACGGAAATCA | *E. coli K12*<br>*Salmonella*<br>*Y. pestis*<br>*Y. pestis KIM* |
| *Binding sequence 5'* yen7 | | | |
| OmpR | 66-79 | TTTGAAATTGAA | *S. enterica* |
| | 72-90 | AATATAATTAGTTTGAAA | *S. enterica* |
| | 103-121 | GTTATTTTTTAATAAAAA | *E. coli* |
| | 129-147 | TAGAAATTATAATGTTAA | *E. coli* |
| PhoB | 22-43 | TTTTTAAATTAATGGTGGTATC | *E. coli* |
| | 68-94 | CTGGCTATAATGCTTAGCACTA | *E. coli* |
| | 129-147 | ATAGCTTAACTGAGAACCT | *E. coli* |

118 Table S5: Primers used in this study

| Primer | Sequence | Temp | Description | Note |
|---|---|---|---|---|
| MS01 | GTGTAGGCTGGAGCTGCTTC | pKD4 | forw primer FRT-Kan-FRT | FRT-site |
| MS02 | CATATGAATATCCTCCTTAGTTCC | pKD4 | rev primer FRT-Kan-FRT | |
| MS101 | AAACATATGACGTTAAAGATTCCGACTG | MH96 | fw pARA-holin(operon) | NdeI |
| MS104 | AAACTCGAGGGATAATGCCGACACACTTTAA | MH96 | rev pARA – hol+m15+lysB | XhoI |
| MS134 | CTAATTGTTCAATAATATGAAC | MH96 | ncRNA220 for | |
| MS135 | GTTAACGTCAGAATGACGCAG | MH96 | ncRNA220 rev | |
| MS136 | ATAGCGATATTGAGCTTATC | MH96 | ncRNA81 for 2 | |
| MS137 | CTTTATAATCAGCTTGAATAGC | MH96 | ncRNA81 rev2 | |
| MS138 | GAGTTTCTCTGTCATTTCGAAG | MH96 | ncRNA673 for | |
| MS139 | CGTGATAAATAGATCTTGGATG | MH96 | ncRNA673 rev | |
| MS29 | CCCTATCTATTAGCTGACCG | MH96 | validation ΔALC fw | |
| MS30 | CACGAACATGTAGAGCCAGC | MH96 | validation ΔALC rev | |
| MS40 | AAACCCGGGCGCGATTGTCTCCTCTTTTTG | MH96 | 1.949 bp 5' *holA* for | SmaI |
| MS41 | AAACCCGGGCCCGATTAAGGAAACTTAG | MH96 | 2.039 bp 3' *rz* rev | SmaI |
| MS42 | AAACCCGGGCCCTCCCCATCCTGTTCTTC | MH96 | 2 bp' 5 *holA* rev | SmaI |
| MS43 | AAACCCGGGAACTACTCCTCCTTAAAATTAG | MH96 | INT for | SmaI |
| MS44 | AAACCCGGGTCATGGGGCAATAGCAGCGC | MH96 | *rz* rev | SmaI |
| MS45 | AAACCCGGGATGATGTATTTTGTAGTGGCAG | MH96 | *roeA* for | SmaI |
| MS46 | AAACCCGGGCCCCCGTTATTTAACGTCAC | MH96 | 3' *rz* for | SmaI |
| MS71 | CAATTACAACGTGGACCTCTGCCG | MH96 | 790 bp 3' *roeA* rev | |
| MS72 | TAAGCGTCGTCCAGTCACCGCCC | MH96 | 1.691 bp 5' *roeA* for | |
| MS74 | AAAGTCGACCATAACTACTCCTCCTTAAAATT | MH96 | lacZ-roeA for | SalI |
| MS75 | AAACCCGGGCCGCAGTTAACGTCAGAATG | MH96 | lacZ-roeA rev | SmaI |
| MS82 | GCGGCGGAGCTCCGGTAGTGGCATAGGGTTAG | MH96 | 2.308 bp 5' *holA* for | SacI |
| MS83 | **GAAGCAGCTCCAGCCTACAC**CCCTCCCCATCCTGTTCTTC | MH96 | 5' region *holA rev* | FRT |
| MS84 | **ACTAAGGAGGATATTCATATG**CCCCCGTTATTTAACGTCAC | MH96 | 3' region *rz* for | FRT |
| MS85 | GCGGCGGAGCTCCCGATTAAGGAAACTTAGTCG | MH96 | 2.039 bp 3' region *rz* rev | SacI |
| MS86 | ATGCGTAAAGGCGAAGAGC | pBAD::sfGFP | sfGFP fw | |
| MS87 | GGATCCTTAATGATGATGATGATGATGTTTGTAC | pBAD::sfGFP | sfGFP rev | Bam HI |
| MS88 | CCCTCTAGAGATTAGATTATGTTCGTTAGCTC | MH96 | Hol 5 fw | XbaI |
| MS89 | **CAGCTCTTCGCCTTTACGCAT**CCCCAAGAAACGCCGCCACAG | MH96 | Hol 5 rev | FRT |
| MS90 | **AAACATCATCATCATCATCATTAA**GGATCCAGTCCGGAGAAAAATCGTGGG | MH96 | Hol 3 fw | FRT Bam HI |
| MS91 | AAACCCGGGCGTTGCTGCAACCGGATAAC | MH96 | Hol 3 rev | XmaI |
| MS92 | AAATCTAGAGGTTCAACAACACGATTCCC | MH96 | Rz1 5 fw | XbaI |

| | | | | |
|---|---|---|---|---|
| MS93 | **CAGCTCTTCGCCTTTACGCAT**TCCTTTTTGGGGAGACATTATT | MH96 | Rz1 5 rev | FRT |
| MS94 | **CAAACATCATCATCATCATCATTAA**GGATCCATCTCTCTTTTAGTCAATTCAG | MH96 | Rz1 3 fw | FRT<br>Bam HI |
| MS95 | AAACCCGGGCCTGAACGCAGTAGGAATC | MH96 | Rz1 3 rev | XmaI |
| MS96 | AAAGGATCCATGCCCGTTCCATACAGAAGC | pHP45 | Spec - BamHI for | Bam HI |
| MS97 | AAAGGATCCACATTATTTGCCGACTACCT | pHP45 | Spec - BamHI rev | Bam HI |
| MS98 | CTGCGTCATTCTGACGTTAAC | MH96 | Validation HolA-sfGFP fw | |
| MS99 | GAGAGATTTATCCTTTTTGG | MH96 | Validation HolA-sfGFP rev | |

Bold letters indicate FRT site, underlined indicate restriction sites

File S1: Differential gene expression H12 vs MH96
